# Deterministic scRNA-seq of individual intestinal organoids reveals new subtypes and coexisting distinct stem cell pools

**DOI:** 10.1101/2020.05.19.103812

**Authors:** Johannes Bues, Marjan Biočanin, Joern Pezoldt, Riccardo Dainese, Antonius Chrisnandy, Saba Rezakhani, Wouter Saelens, Vincent Gardeux, Revant Gupta, Julie Russeil, Yvan Saeys, Esther Amstad, Manfred Claassen, Matthias Lutolf, Bart Deplancke

## Abstract

Single-cell RNA-sequencing (scRNA-seq) has transformed our ability to resolve cellular properties across systems. However, current scRNA-seq platforms are one-size-fits-all approaches that are tailored toward large cell inputs (> 1,000 cells), rendering them inefficient and costly when processing small, individual tissue samples. This important drawback tends to be resolved by loading bulk samples, but this yields confounded mosaic cell population read-outs. To overcome these technological limitations, we developed a <u>d</u>etermin<u>is</u>tic, mRNA-capture bead and cell <u>co</u>-encapsulation dropleting system, DisCo. We demonstrate that DisCo enables precise particle and cell positioning and droplet sorting control through combined machine-vision and multilayer microfluidics. In comparison to other microfluidics systems, the active flow control driving DisCo, enables continuous operation and processing of low-input samples (< 100 cells) at high capture efficiency (> 70%). To underscore the unique capabilities of our approach, we analyzed intestinal organoid development by “DisCo-ing” 31 individual organoids at varying developmental stages. This revealed extensive organoid heterogeneity, identifying distinct subtypes including a regenerative fetal-like *Ly6a*^+^ stem cell population which persists as symmetrical cysts even under differentiation conditions. Furthermore, we uncovered a so far uncharacterized “gobloid” subtype consisting predominantly of precursor and mature (*Muc*2^+^) goblet cells. These findings demonstrate the unique power of DisCo in providing high-resolution snapshots of cellular heterogeneity among small, individual tissues.

## Introduction

Single-cell RNA sequencing (scRNA-seq)^1^ induced a paradigm shift in biomedical sciences, since it allows the dissection of cellular heterogeneity by high-dimensional data. Recent technological developments, particularly for cell capture and reaction compartmentalization^2–6^, have led to a substantial increase in experimental throughput, enabling massive mapping efforts such as the mouse and human cell-atlas studies^5,7,8^. These developments were accompanied by biochemical advances, for instance for targeted transcript detection or library multiplexing^9,10^, which present a rich toolbox for large-scale scRNA-seq studies. However, since the majority of methods rely on stochastic cell capture, entailing large sample inputs, efficient processing of small samples (< 1,000 cells) remains challenging. The three main reasons for this are: 1) high fixed run costs, which lead to a large expense per cell at low inputs. For instance, a 10X Chromium run on 100 cells would cost $44 per sequenced cell. 2) Requirements of minimum cell inputs. For example index-sorting FACS or 10X Chromium require minimum cellular inputs ranging between 10,000 and 500 cells, respectively^11,12^. 3) Reduced effectiveness at low inputs because of limited cell capture efficiencies or cell size-selective biases^13^ when processing small heterogeneous samples. To illustrate these limitations, we summarized the performance of various scRNA-seq technologies on low input samples in **Supplementary Table 1**. Consequently, small samples, involving for instance zebrafish embryos^14^, organisms like *C. elegans*^15^, or intestinal organoids^16–18^, are still pooled to obtain cell numbers that are compatible with stochastic microfluidic and well-based technologies. Thus, it is rather paradoxical that limitations overcome by single cell methods are nevertheless reintroduced at the sample level: artificial averages across samples, resulting in an inability to resolve cell type distributions of individual systems or tissues. This particularly hampers research on emergent and self-organizing multicellular systems, such as organoids, that are heterogeneous and small at critical development stages.

In this study, we develop a novel <u>d</u>etermin<u>is</u>tic, mRNA-capture bead and cell <u>co</u>-encapsulation dropleting system (DisCo) for low input scRNA-seq. In contrast to established methods that rely on passive cell capture strategies, we utilize machine-vision to actively detect cells and coordinate their capture in droplets. This active flow control approach allows for continuous operation, enabling free per run scaling and serial processing of samples. We demonstrate that DisCo can efficiently process samples of 100 cells and below, making this platform well suited to handle small, individual tissues. Here, we exploit DisCo’s unique capabilities to explore the heterogeneous early development of single intestinal organoids at the single cell level. Grown from single stem cells, organoids of vastly different morphologies and cell type compositions form under seemingly identical *in vitro* conditions^16^. These unpredictable developmental patterns represent one of the major limitations of this model system, preventing their widespread implementation e.g. in drug screens^19^. Thus, efforts to advance our understanding of the extent of organoid heterogeneity, how it arises, and how it can be controlled, for instance with synthetic growth matrices^20,21^, are of essence. In depth mapping of individual organoid heterogeneity by scRNA-seq has so far been prevented by the minute cell numbers contained in a single intestinal organoid at critical developmental stages, such as post symmetry breaking at the 16-32 cell stage^16^. In total, we “DisCo’d” 31 single organoids at four developmental time points post symmetry breaking, and identified striking differences in cell type composition between individual organoids. Among these subtypes, we detected “spheroids” that are composed of regenerative fetal-like stem cells marked by Stem Cell Antigen-1 (*Sca1*/*Ly6a*)^22–25^ and that persist under differentiation conditions. In addition, we uncovered a rare subtype that is predominantly comprised of precursor- and mature goblet cells, which we term “gobloids”.

## Results

To develop our Deterministic Co-encapsulation (DisCo) system, we engineered a three inlet (cells, beads, oil) multilayer dropleting device with two outlet ports (sample, waste) (Schematic **Figure 1A**, full design **Supplementary Figure 1A**). On this device, each inlet and outlet was augmented with a Quake-style microvalve^26^, to facilitate flow control during operation. In addition, one common valve spanning both the cell and bead channel, termed the dropleting valve, was integrated to allow for on-demand droplet generation. To operate the device, we developed a three-stage process (**Figure 1B**): 1. Stop two particles at the encapsulation site, 2. Eject particles into one droplet, 3. Selectively extract the droplet in a sample channel (Microscopy images of the process are depicted in **Figure 1C**). To enable precise coordination of particles in microchannels, we developed a machine-vision-based approach utilizing subsequent image subtraction for blob detection (**Supplementary Figure 1B**), and on-chip valves for flow-control. Deterministic displacement patterns were induced by opening and closing the cell and bead valves (depicted in **Supplementary Figure 1C)**, which moved particles according to discrete jumps into the target region of interest (ROI) with 95.9% of particles placed in an approximately ~200 μm wide region (**Supplementary Figure 1D**). Upon placement, the stopped particles were ejected by pressurizing the dropleting valve, displacing an equal volume of liquid from both channels. The ejected liquid phase was then sheared into a droplet by activating the oil stream. We found that precise pressurization of the dropleting valve allowed for accurate control of droplet volume (**Supplementary Figure 1E**, **Supplementary Video 1**). Post droplet formation, the outlet valves were actuated to separate the formed droplet from the excess waste liquids (**Figure 1D**). With all components operating in tight orchestration, we were able to generate monodisperse emulsions with high co-encapsulation purity (**Figure 1E**, **Supplementary Video 2**).

**Figure 1.**
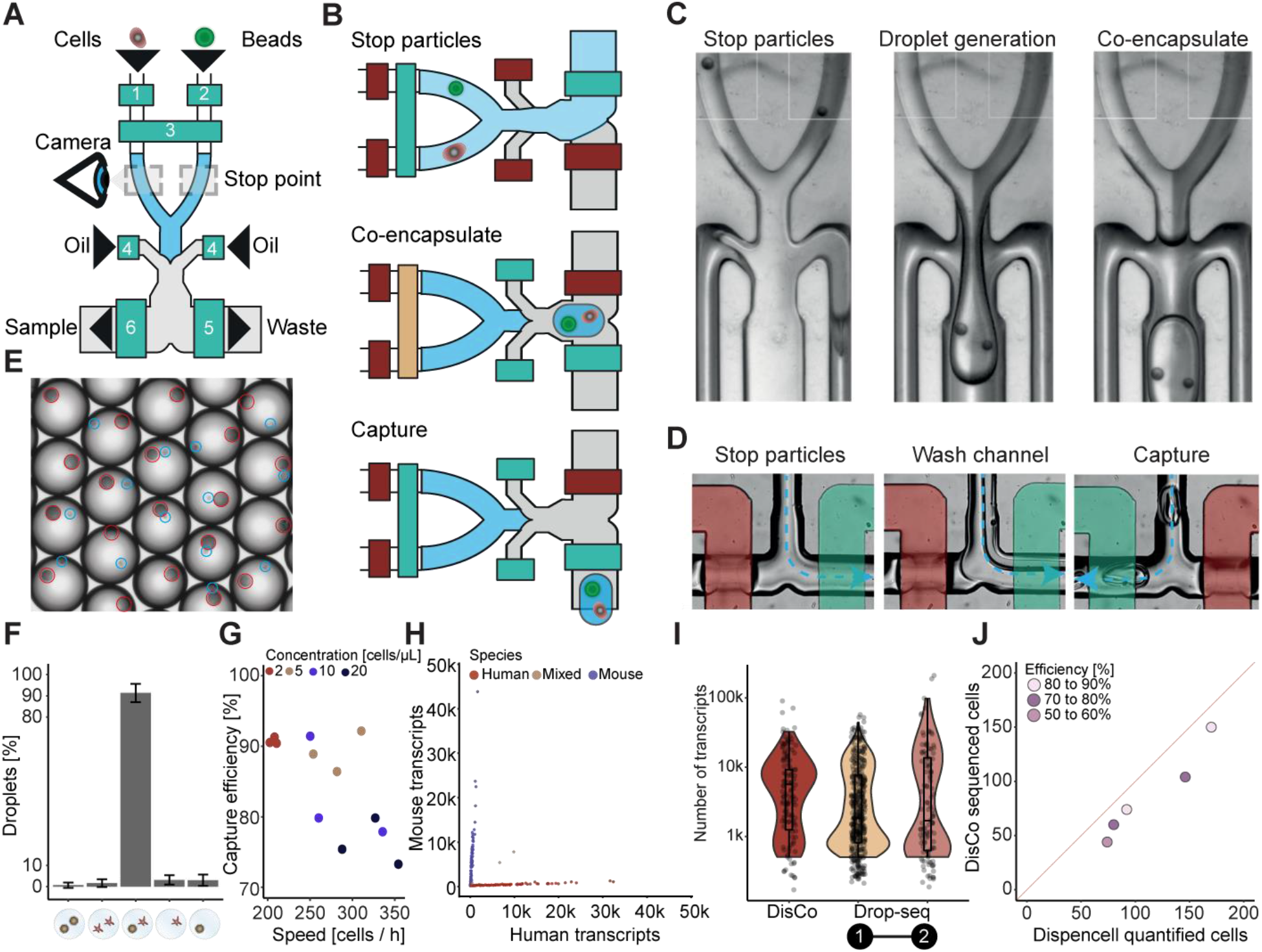
Overview and critical feature assessment of the <u>d</u>etermin<u>is</u>tic <u>co</u>-encapsulation (DisCo) system: (**A**) Schematics of the DisCo microfluidic device. The device contains three inlet channels for cells, beads, and oil, and two outlets for waste and sample liquids. All inlets and outlets are augmented with Quake-style microvalves (green boxes): 1. cell valve, 2. bead valve, 3. dropleting valve, 4. oil valve, 5. waste valve, 6. sample valve. The device is continuously monitored by a high-speed microscopy camera to detect and coordinate placement of particles at the Stop point. (**B**) Illustration of the particle co-encapsulation process on the DisCo device. Initially, two particles (here a bead and a cell) are stopped (Stop particles) in close proximity to the channel junctions by closing the channel valves (red: closed, green: open). Next, by pressurizing the dropleting valve (yellow), both particles are ejected into the junction point, and the droplet is sheared by opening the oil valve (Co-encapsulate). Finally, the produced droplet is captured in the Sample channel (Capture). (**C**) The co-encapsulation process of two beads and droplet generation as observed on chip. Dyed liquids were used to examine the liquid interface of the carrier liquids. Channel sections with white squares are 100 μm wide. (**D**) The droplet capture process as observed on-chip. Valves are highlighted according to their actuation state (red: closed, green: open). While particles are stopped, excess buffers are discarded through the waste channel and the channel is flushed with oil prior to droplet capture. Upon co-encapsulation, the waste valve is closed, the sample valve opened, and the produced droplet captured in the Sample channel. (**E**) Images of DisCo droplet contents. Cells (blue circle) and beads (red circle) were co-encapsulated, and captured droplets imaged. Mean bead-size is approximately 30 μm. (**F**) Droplet occupancy of DisCo-processed cells and beads for cell concentrations ranging from 2 to 20 cells per μl (total encapsulations n = 1203). Error bars represent standard deviation. (**G**) Cell capture efficiency and cell capture speed for varying cell concentrations (total encapsulations n = 1203). Cells were co-encapsulated with beads at concentrations ranging from 2 - 20 cells per μl, and co-encapsulation events quantified by analyzing recordings of the process. (**H**) DisCo scRNA-seq species separation experiment. HEK 293T and murine pre-adipocyte iBA cells were processed with the DisCo workflow for scRNA-seq, barcodes merged, and species separation visualized as a Barnyard plot. (**I**) Comparison of detected UMIs per cell of conventional Drop-seq experiments. UMIs per cell from HEK 293T data for conventional Drop-seq experiments ([1] - from Biočanin, Bues *et al.* 2019^27^ and [2] - from Macosko *et al.* 2015^2^), compared to the barcode-merged HEK 293T DisCo data. Drop-seq datasets were down-sampled to comparable sequencing depth. Box elements are described in the **Materials and Methods** section. (**J**) Total cell processing efficiency of DisCo at low cell inputs. Input cells (HEK 293T) ranging from 74 to 170 were quantified with the Dispencell system. Subsequently, all cells were processed with DisCo, sequenced, and quality filtered (> 500 UMIs). The red line represents 100% efficiency, and samples were colored according to recovery efficiency after sequencing.

As a first benchmarking experiment, we set out to determine the encapsulation performance of DisCo for scRNA-seq-related applications, involving co-encapsulation of single cells with microspheres. Specifically, we aimed to reconfigure the Drop-seq^2^ approach as it only requires coordination of two channels, as compared to three channels for inDrop^3^. Since co-encapsulation purity and cell capture efficiency are critical system parameters for droplet scRNA-seq systems, we quantified the system’s processing speed and encapsulation performance in a free-run configuration, i.e. without cell number limitations at varying cell densities. We found that on average, 91.4% of all droplets contain a cell and a bead, and 1.7% contain an independent cell doublet (**Figure 1F**). Overall, the system provided high cell capture efficiencies of 90% at around 200 cells per hour for a 2 cells/μL cell concentration (**Figure 1G**). At higher cell concentrations of 20 cells/μL, the processing speed could be increased to 350 cells per hour, yet with decreased capture efficiencies of approximately 75%.

Next, we benchmarked the performance of DisCo for scRNA-seq. With drastically reduced bead amounts contained in the generated sample emulsion, we utilized a previously developed chip-based cDNA generation protocol^27^. Initially, as a library quality measure, we performed a species-mixing experiment of human HEK 293T and murine brown pre-adipocyte IBA cells. We observed clear species separation (**Figure 1H**), consistent with the limited number of previously detected doublets (**Figure 1F**), and increased read-utilization rate compared to conventional Drop-seq experiments (**Supplementary Figure 1F**). As previously reported^28^, we found that our data displayed a skewed barcode sequence editing distance distribution compared to a true random distribution (**Supplementary Figure 1G**). Since the uniquely low number of beads in DisCo samples (< 500) renders the random occurrence of barcode sequences with an editing distance < 3 rare, we developed a graph-based approach to identify and merge closely related barcodes (described in **Material and Methods**). We found that this approach did not compromise the single cell purity (**Supplementary Figure 1H**) and improved the detectable number of transcripts per cell as compared to published Drop-seq datasets on HEK 293T cells^2,27^ (**Figure 1I**).

Since DisCo actively controls fluid flow on the microfluidic device, we observed that the system requires negligible run-in time, and is capable of efficiently processing cells from the first cell on. Given this observation, and the high-capture efficiency of DisCo in free-run mode, we hypothesized that the system should provide reliable performance on small samples of 100 cells and below. To determine the overall cell capture efficiency of DisCo, we precisely quantified the number of input cells using impedance measurements. Specifically, we utilized custom pipette tips augmented with a DISPENCELL gold-plated electrode, which allowed accurate counting of the number of input cells as validated by microscopy (**Supplementary Figure 1I**). Utilizing the DISPENCELL approach, we processed cell numbers between 50 - 200 cells, of which on average 86.4% (SD ± 8.1%) were visible on the chip. Of all input cells, 79.1% (SD ± 7.4%) were successfully co-encapsulated, which corresponds to a co-encapsulation efficiency of 91.6% (SD ± 1.6%) of all visible cells, while 74.9% (SD ± 10.7%) of input cells were found as barcodes over 500 UMIs per cell (**Figure 1J**).

As a real-world application, we used DisCo to explore the developmental heterogeneity of intestinal organoids^29^. These polarized epithelial tissues are generated by intestinal stem cells in 3D matrices through a stochastic self-organization process, and mimic key geometric, architectural and cellular hallmarks of the adult intestinal mucosa (e.g. a striking crypt-villus-like axis)^29^. When grown from single stem cells, organoids of very different morphologies form under seemingly identical *in vitro* conditions (**Figure 2A,** overview image in **Supplementary Figure 2A**). Pooled tissue scRNA-seq data has shed light on the *in vivo*-like cell type composition of these organoids^16–18,30^, but cannot resolve inter-organoid heterogeneity. Critical for organoid development is an early symmetry breaking event at Day 2 (16-32 cell stage) that is triggered by cell-to-cell variability and results in the generation of the first Paneth cell responsible for crypt formation^16^. Here, we were particularly interested in examining the emergence of heterogeneity between individual organoids subsequent to the symmetry breaking timepoints. To do so, we isolated single LGR5^+^ cells by FACS, and maintained them in a stem cell state using CHIR99021 and valproic acid (CV)^31^. On Day 3 of culture, CV was removed to induce differentiation. In total, we sampled 31 single intestinal organoids across four timepoints (Day 3 - 6) (**Figure 2A**). These organoids were selected based on differences in morphology, and may thus not constitute an unbiased sample of the population. Since Day 3 represents both differentiation Day 0 and the first sampling time point, we re-annotated the data accordingly (S0 – S3 replacing Day 3 – Day 6). During the co-encapsulation run, the number of encapsulated cells was noted and correlated to the number of barcodes retrieved, which was in approximate accordance (**Supplementary Figure 2B**). The even distribution of the number of reads mapping to ribosomal protein transcripts and the observed low expression of heat shock protein-coding genes indicates that most cells were not affected by dissociation and on-chip processing (**Supplementary Figure 2C**).

**Figure 2.**
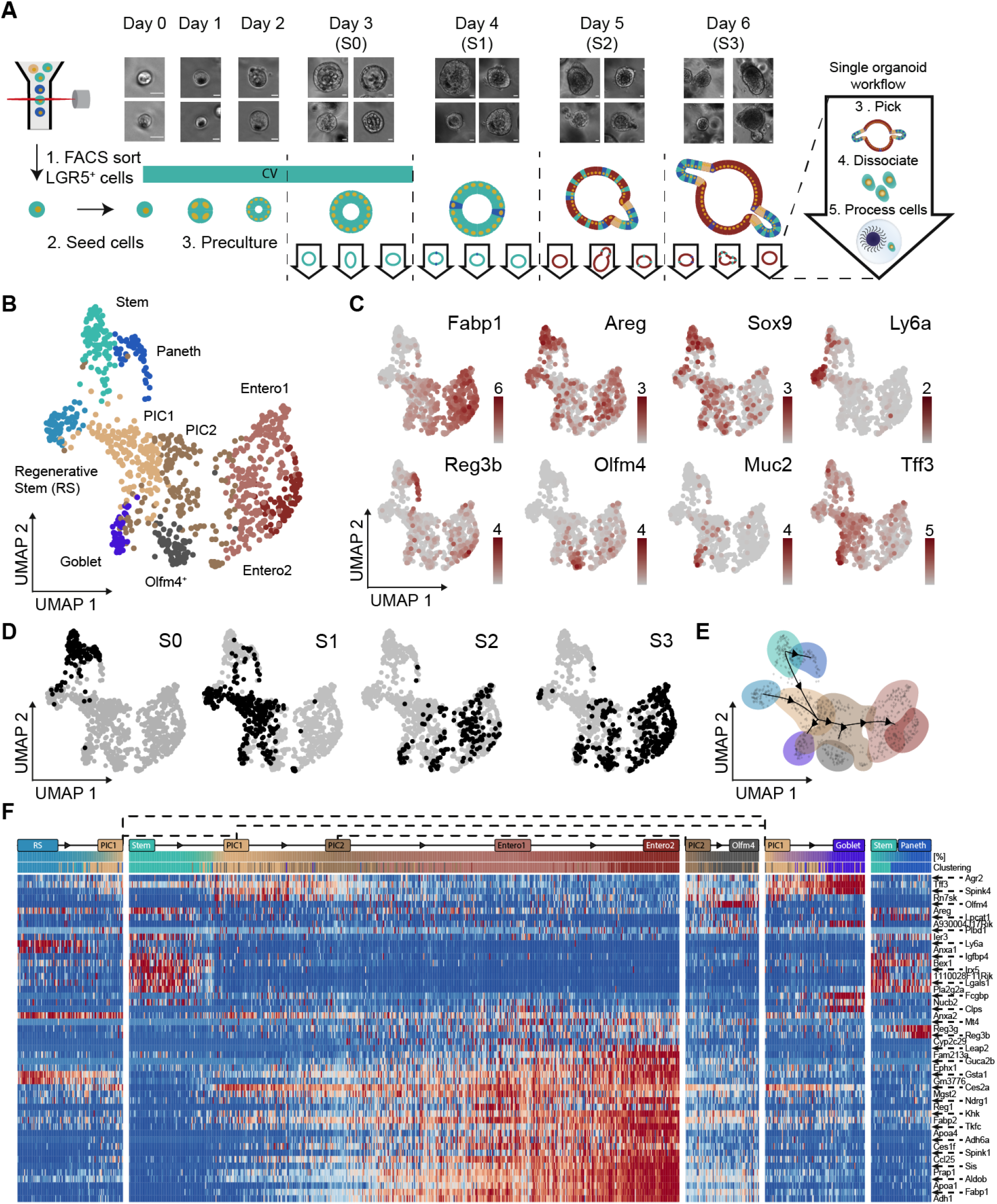
Utilizing DisCo to map intestinal organoid cell heterogeneity along development: (**A**) Overview of the experimental design for DisCo’ing individual organoids. Single LGR5+ intestinal stem cells were isolated via FACS and precultured for 3 days under stem cell maintenance conditions (ENR CV Day 0 to 3). On Day 3, CV was removed from the culture, and organoids differentiated under ENR conditions for up to 3 days. For each day during development (S0 - S3), individual organoids were isolated, dissociated, and processed on the DisCo platform. Representative bright-field imaging examples of individual organoids for each day are shown on top. Scale bar 50 μm. (**B**) UMAP embedding of all sequenced cells. All 945 processed cells from 31 organoids were clustered with k-means clustering, after which clusters were annotated according to marker gene expression. (**C**) UMAP-based visualization of the expression of specific markers that were used for cluster annotation. (**D**) Temporal occurrence of cells. Cells are highlighted on the UMAP embedding according to sampling time point (S0 - S3). (**E**) Developmental trajectory based on the cluster annotation and the sampling time point derived by slingshot^34^. Cells were annotated in accordance with clustering in (B). (**F**) Heat map of differentially expressed genes along the waypoints of the trajectory. Waypoints are annotated in accordance with cell clustering as in (B). Cluster abbreviations: Stem cells (Stem), Regenerative stem cells (RS), Potential intermediate cells (PIC)^33^, Enterocytes cluster 1/2 (Entero1/2).

To retrieve a first overview of overall cellular heterogeneity, we jointly visualized all 945 cells passing the quality thresholds through Uniform Manifold Approximation and Projection (UMAP). We found that our data was consistent with previously published pooled organoid scRNA-seq read-outs^17,30^ since it revealed expected cell types including *Fabp1*-expressing enterocytes, *Muc2*-expressing goblet cells, *Reg3b*-positive Paneth cells, and *Olfm4*-expressing stem cells (**Figure 2B** and **2C**). In addition, a rare subset of cells, likely too few to form clusters, showed *ChgA* and *ChgB* expression, indicating the expected presence of enteroendocrine cells (**Supplementary Figure 2D**). Noteworthy, we found that batch effects are correctable since no batch-based clustering was observed after correction (**Supplementary Figure 2E**). We also did not detect any clustering driven by cell quality, e.g. detected transcripts or mitochondrial transcripts (**Supplementary Figure 2C**). These findings support the cell type-resolving power of our DisCo platform (**Figure 2C,** extensive heatmap in **Supplementary Figure 2F**, and list in **Supplementary Table 2**). In addition to the expected cell types, we observed a distinct cluster marked by high expression of Stem cell antigen 1 (*Sca1* or *Ly6a*). In depth analysis of marker genes showed high expression of *Anxa1* and *Clu* in the same cluster (**Supplementary Figure 2D**), and increased YAP-1 target gene expression (**Supplementary Figure 2G**), suggesting that these cells are most likely regenerative fetal-like stem cells^24,25,32^. Since the two remaining clusters did not show a striking marker gene signature, we resolved their identity by imposing temporal information on the data. This revealed that these clusters likely represent stem- and previously termed potentially intermediate cells (PIC)^33^, given their occurrence at early developmental time points (**Figure 2D**). As expected, mature cell types were mostly present at later time points. To further leverage the temporal component in the DisCo data, we used slingshot trajectory analysis^34^ to infer lineage relationships between cell types and to identify genes that may be of particular significance for waypoints along differentiation (**Figure 2E**). Beyond the previously utilized marker genes for cell type annotation, for example *Reg3b* and *Reg3g* for Paneth cells, additional established markers^35^ were identified, such as *Agr2 and Spink4*, and *Fcgbp* for goblet cells (**Figure 2F**). Overall, this suggests that the meta-data produced with our DisCo platform aligns with and expands prior knowledge.

Intriguingly, we observed maintained presence of the *Ly6a*+ stem-cell population at S0, S1, and S3. Since cells with similar expression signatures were previously described under alternate culture conditions as belonging to a distinct organoid subtype termed spheroids^23^, we next aimed to verify the presence of such spheroids among our sampled organoids and study their temporal behavior. To do so, we stratified our cells according to the individual organoids from which they were derived by mapping this information onto the reference scaffold (**Figure 3A**). Globally, this analysis revealed that the maturation seems to follow the expected pattern with early organoids (S0) mainly containing stem and Paneth cells, and older organoids (S1 – S3) differentiated cells like goblet cells and enterocytes. However, within single organoids, we found strong heterogeneity, revealing that *Ly6a*^+^ cells were indeed present in a distinct subset of organoids, predominantly comprised of these cells (S1a, S3e). Furthermore, images obtained prior to dissociation showed that *Ly6a*^+^ cell-containing organoids (S3e) exhibited a larger, cystic like structure (**Supplementary Figure 3A**). To confirm the presence of *Ly6a*^+^ organoids in our cultures, we utilized RNAscope (**Figure 3B,** controls **Supplementary Figure 3B**) to localize *Ly6a*, *Muc2,* and *Fabp1* expression in organoid sections. These analyses revealed canonical budding organoids, containing few *Muc2*^+^ goblet cells and *Fabp1*^+^ enterocytes, and *Ly6a*-expressing cells in spherical organoids that did not contain differentiated cell types such as enterocytes or goblet cells.

**Figure 3.**
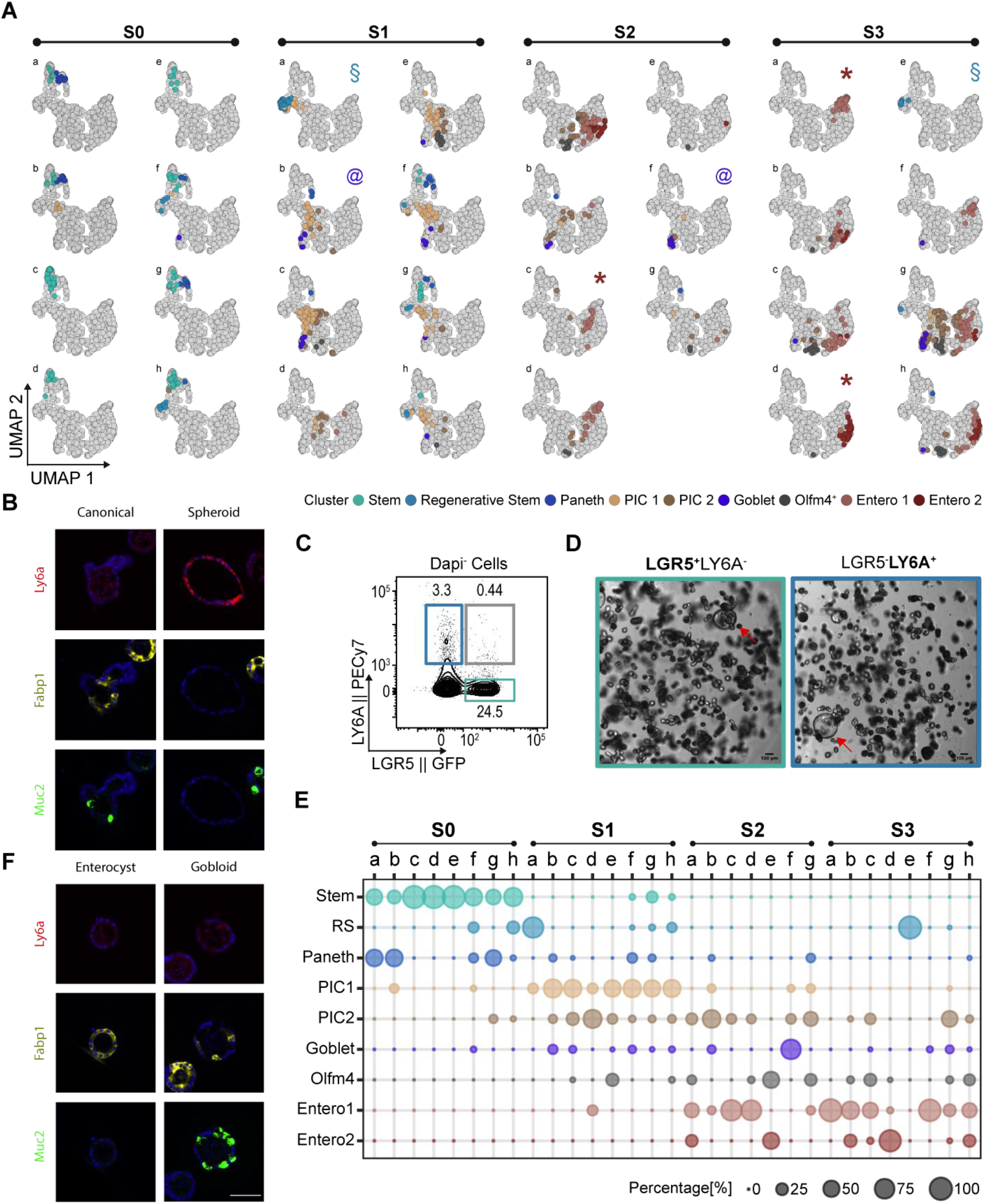
Cell type distribution and marker gene expression across individual intestinal organoids during development: (**A**) Projection of cell types onto 31 individual organoids. Cells per single organoid were colored according to their global clustering and highlighted on the UMAP embedding of all sequenced cells. Projections are grouped according to their sampling time. Manually classified organoids were annotated with the following symbols: “*” enterocysts, “§” spheroids, “@” gobloids. **(B)** *in situ* RNA detection of *Ly6a*, *Fabp1*, and *Muc2* expression. A representative canonical and *Ly6a*-expressing organoid is displayed. Scale bar (displayed in F) 50 μm. (**C**) Surface LY6A and LGR5-GFP expression under ENR CV conditions. The dot plot depicts LGR5-GFP and LY6A expression in organoid-derived single cell suspensions. The numbers indicate frequencies (%). (**D**) Culturing outcomes of LGR5+cells and LY6A^+^ cells. Single LGR5^+^ LY6A^−^ and LGR5-LY6A^+^ cells were isolated by FACS and seeded in Matrigel. Cells were cultured as depicted in **Figure 2a** and imaged using bright-field microscopy at S3. Red arrows point to spheroid morphologies. Scale bar 100 μm. (**E**) Dotplot depicting the distribution of annotated cell types per organoid. Dot size depicts the percentage of cells associated to each cluster per organoid. **(F)** *in situ* RNA detection of *Fabp1* and *Muc2* expression. Selected images resembling the enterocyst and gobloid subtypes. Scale bar 50 μm.

The presence of *Ly6a^+^* cells during the first day of sampling suggested that these cells constitute a second, *Lgr5*-independent stem cell population in the organoid culture. Using flow cytometry, we found that the majority of cells are either LGR5^+^ LY6A^−^ (24.5 %) or LGR5^−^ LY6A^+^ (3.3 %) with only a minority (0.4%) being double positive (**Figure 3C**). This finding, in combination with our trajectory analysis (**Figure 2E** and **2F**), suggested that *Ly6a*^+^ cells are capable of differentiating into organoids. To test this, we sorted and differentiated LGR5^−^ LY6A^+^ cells, revealing that both LGR5^+^ LY6A^−^ and LGR5^−^ LY6A^+^ cells give rise to organoids of similar morphological heterogeneity (**Figure 3D**). These results indicate that LGR5^−^ LY6A^+^ cells have full stem cell potential, comparable to that of previously described fetal-like stem cells^23^. Furthermore, the fact that LGR5^−^ LY6A^+^ cells did not display a propensity towards spheroid formation suggests that environmental conditions, e.g. matrix stiffness, rather than the initial cell state dictate the formation of spheroids.

Beside the *Ly6a*^+^ cell-enriched organoids, our data suggested the presence of additional organoid subtypes in the per organoid mappings (**Figure 3A**). The two most striking additional subtypes were three organoids that contained mostly enterocytes (S2c, S3a, S3d), and two that consisted predominantly of immature and mature goblet cells (S1b and especially S2f). The identity of the observed subtypes was further substantiated when visualizing the cell type abundance per organoid (**Figure 3E**), and marker gene expression in individual organoids (**Supplementary Figure 3C**). Similar to the spheroids, both subtypes showed aberrant morphologies, tending to be small and round, as compared to canonical organoids bearing a crypt-villus axis (e.g. S3c, **Supplementary Figure 3A**). To detect more subtle molecular differences, we used psupertime^36^ to identify genes that are dynamically expressed during the development of individual organoids. This analysis revealed additional genes that are expressed in subsets of organoids, such as Gastric inhibitory polypeptide (*Gip*), Zymogen granule protein 16 (*Zg16*), Vanin 1 (*Vnn1*), and Defensin alpha 24 (*Defa24*) (**Supplementary Figure 3D**).

While organoids dominated by enterocytes were previously described as enterocysts^16^, organoids displaying goblet cell hyperplasia, here termed “gobloids”, were so far to our knowledge unknown. To validate the existence of the uncovered organoid subtypes, we utilized RNAscope to localize the expression of enterocyte (*Fabp1*) and goblet cell (*Muc2*) markers (**Figure 3F,** controls in **Supplementary Figure 3B**). In addition, and in agreement with our data and prior research, we detected organoids that exclusively contained *Fabp1*^+^ cells, most likely representing enterocysts. Most importantly, we were able to identify organoids that contained a high number of *Muc2^+^* goblet cells, confirming the existence of “gobloids”.

## Discussion

A key feature of our new DisCo approach is the ability to deterministically control the cell capture process. Despite lowering the throughput compared to stochastic droplet systems^2,3^, our approach provides the advantage of being able to process low cell input samples at high efficiency and at a strongly decreased per cell cost (**Supplementary Table 1**). Thus, we believe that the DisCo approach is filling an important gap in the scRNA-seq toolbox. Moreover, full control over the encapsulation process allows for continuous operation of our platform, which is offsetting to some extent the decreased throughput. Another critical feature of DisCo is the use of machine-vision to obtain full control of the entire co-encapsulation process including particle detection, particle positioning, particle droplet injection, and droplet volume. This enables the correct assembly of most droplets, virtually eradicating confounding factors that arise due to failed co-encapsulations^37,38^. In concept, DisCo is thus fundamentally different to passive particle pairing approaches such as traps^39–41^ and, compared to these technologies, offers the advantage of requiring vastly simpler and reusable chips without suffering from cell/particle size and shape selection biases^13,42^. This renders the DisCo approach universally applicable to any particle co-encapsulation application^43,44^, i.e. cell-cell encapsulations, with the only limiting factor being particle visibility. Providing further development, we envision that machine learning-based deterministic cell handling will ultimately enable targeted cell selection, e.g. by fluorescence or morphology, transforming DisCo into an end-to-end cell processor for samples with low-to-medium input samples.

To demonstrate DisCo’s capacity to process small tissues/systems that were so far difficult to access experimentally, we have analyzed the cell heterogeneity of chemosensory organs from *Drosophila* larvae^45^ and, as shown here, single intestinal organoids. It is thereby worth noting that, based on our handling of distinct tissues, we found that not DisCo itself, but rather cell dissociation has become the efficiency-limiting factor, a well-recognized challenge in the field^46,47^. Indeed, substantial cell loss was a regular occurrence, even with optimized dissociation and processing strategies (see **Methods**).

scRNA-seq of individual organoids led us to uncover organoid subtypes of aberrant cell type distribution that were previously not resolved with pooled organoid scRNA-seq^16,17,30^. One subtype contained predominantly cells that were strikingly similar to previously described fetal-like stem cells or revival stem cells that occur during intestinal regeneration^24,25,32^. This subtype, previously described under alternate culture conditions as spheroid-type organoids^18,22,23^, was identified here under standard organoid differentiation conditions, indicating that these organoids are capable of maintaining their unique state. We isolated LY6A-expressing cells and found that they readily give rise to canonical organoids, indicating that these cells are capable of providing a pool of multipotent stem-cells. Of particular interest was one organoid subtype that we termed “gobloid” given that it predominantly comprises immature and mature goblet cells. Since low Notch signaling is pivotal for the commitment of crypt base columnar (CBC) cells towards secreting progenitors, lack of Notch ligand-providing Paneth cells^48^, may drive gobloid development^49^. However, failure to produce Paneth cells has previously been suggested as a mechanism underlying enterocyst development^16^, which in principle requires high Notch signaling. Hence, we believe that our findings establish an important foundation to support further research on the emergence of gobloids and enterocysts from the still elusive PIC cells, providing an exciting opportunity to delineate lineage commitment factors of CBC cell differentiation.

In sum, we demonstrate that our DisCo analysis of individual organoids is a powerful approach to explore tissue heterogeneity and to yield new insights into how this heterogeneity arises. In comparison to established approaches such as automated microscopy^16,18^, DisCo is magnitudes lower in experimental scale. Nevertheless, scRNA-seq data acquired from 31 organoids enabled us to recapitulate previous findings, benchmarking DisCo, and most importantly, to uncover novel subtypes, leveraging the key advantage of scRNA-seq, i.e. independence from *a priori* knowledge. Next to catalyzing research on other tissues or systems of interest, we believe that the technology and findings of this study will contribute to future research on intestinal organoid development and thus aid the engineering of more robust organoid systems. Furthermore, we expect this approach to be applicable to rare, small clinical samples to gain detailed insights into disease-related cellular heterogeneity and dynamics.

## Acknowledgements

We thank Wanze Chen and Petra C. Schwallie for constructive discussions. We thank Virginie Braman for help in establishing intestinal organoid culture in our group, and Giovanni Sorrentino from Kristina Schoonjans’ lab for valuable advice and support during organoid culture establishment. We also thank Luc Aeberli and Georges Muller from SEED Biosciences for cell sorting support. We thank the EPFL CMi, GECF, BIOP, FCCF, Histology core facility, SCITAS, and UNIL VITAL-IT for device fabrication, sequencing, imaging, sorting, histology, and computational support respectively. We particularly thank Jessica Sordet-Dessimoz for her support with the RNA-scope assay. This research was supported by an Animalfree Research 3R Grant, the Swiss National Science Foundation Grant (IZLIZ3_156815) and a Precision Health & related Technologies (PHRT-502) grant to B. D., the Swiss National Science Foundation SPARK initiative (CRSK-3_190627) and the EuroTech PostDoc programme co-funded by the European Commission under its framework programme Horizon 2020 (754462) to J. P., as well as by the EPFL SV Interdisciplinary PhD Funding Program to B. D. and E. A.. Y.S. is an ISAC Marylou Ingram scholar.

## Contributions

BD, JB, and MB designed the study. BD, JB, MB, and JP wrote the manuscript. JB and RD designed and fabricated microfluidic chip. JB developed the machine-vision integration for DisCo. JB and MB benchmarked the system and performed all single-cell RNA-seq experiments. JP, JB, MB, WS, VG, and RG performed data analysis related to single organoid scRNA-seq experiments. JB, SR and MB performed all organoid and cell culture assays. JB, AC, and JR, performed all imaging assays. EA provided critical comments regarding microfluidic chip design and fabrication. MC provided critical comments on intestinal organoid scRNA-seq data analysis. ML provided critical comments regarding intestinal organoid scRNA-seq data and design of critical confirmation experiments. All authors read, discussed, and approved the final manuscript.

