## Supplementary materials for "Deterministic scRNA-seq of individual intestinal organoids reveals new subtypes and coexisting distinct stem cell pools"

1. Materials and Methods
2. Supplementary Figures 1-3
3. Supplementary Tables 1 and 2

### 1. Materials and Methods

##### System comparison metrics

Performance metrics for (Supplementary Table 1) were calculated the following ways:

- **Minimum cell input estimates:** The minimum cell input values were derived from the following sources: **10X Chromium**[1]**:** Lowest cell input number from the 10X Chromium manual (CG000183 Rev C); **inDrop**[2]**:** Lowest numbers mentioned in the 1CellBio manual (Single Cell Encapsulation Protocol, Version 2.4); **Drop-seq**[3]**:** Lowest numbers utilized in Zhang et al. 2019[4]. It is likely that lower cell numbers can be processed, yet Drop-seq has been suggested to be used “When the sample is abundant” by Zhang et al. 2019; **FACS based methods**[5,6]**:** Input limits as described by Hwang et al. 2018[7]; **Fluidigm C1**[8]**:** Lowest cell input number from the Fluidigm C1 specification sheet (Specification Sheet PN 101-3387 D1); **Wafergen iCell8**[9]**:** Lowest cell numbers were derived from the iCell manual (CELL8 Single-Cell ProtocolD07-000025 Rev. C). According to the manual, 80 µL of 0.02 cells/nL suspension are prepared for dispensing; **Seq-well**[10]**:** The lowest cell number used for capture in Gierahn et al. 2017[10]. **Disco:** The lowest cell number processed in this study.
- **Efficiency estimates:** Efficiency estimates were derived from varying sources and represent different efficiencies. The efficiencies for **10X Chromium**, **inDrop**, and **Drop-seq** were derived from Zhang et al. 2019[4] from quantified cellular inputs (> 1000 cells) and sequenced cells passing quality thresholds. Since these efficiencies stem from experiments that were performed with optimized cell inputs, we can assume lower efficiencies when processing low cell inputs (< 1000). The efficiency for the **Fluidigm C1** system was derived from Xin et al. 2016[11] from a high input sample of primary cells. The efficiency represents a conversion efficiency from captured to sequenced cells passing quality thresholds, thus it does not include cell capture inefficiencies which are substantial at low cell inputs (personal communication Dr. Bastien Mangeat, Gene Expression Core Facility EPFL). For the **Wafergen iCell8** system, an efficiency estimate was derived from Wang et al. 2019[12] and represents the conversion efficiency from captured to sequenced cells passing quality thresholds, thus it does not include cell capture inefficiencies. The efficiency for **Seq-well** was derived from Gierahn et al. 2017[10] at 400 cells input and represents an inferred

efficiency from quantified cell input to sequenced cells passing quality thresholds. Specifically, the library conversion efficiency, i.e. the percent of captured cells identified in the sequencing data passing quality thresholds, was calculated based on the species-mixing experiment involving 10,000 input cells. The library conversion efficiency, in combination with capture efficiencies at 400 cells, was utilized to determine the efficiency at low cell numbers. Hence, this is inferred from quantified cellular inputs to sequenced cells passing quality thresholds. **DisCo:** The efficiencies were derived in this study and represent mean efficiencies for low cell inputs (50 - 200), from quantified cell input to sequenced cells passing quality thresholds.

- **Cost per cell estimates:** Two cost estimate numbers are listed for 100 cells i) the cost for 100 cells not considering system efficiencies ($/cell, 100 output cells), and ii) the cost for 100 input cells considering the listed efficiencies ($/cell, 100 input cells). Run costs for **Smart-seq2**, **Cel-seq2**, **inDrop**, **Drop-seq**, and **Seq-well** were derived from Ding et al. 2020 (Supplementary Table 8)[13]. Run costs for **10X Chromium**, **Fluidigm C1** (96), and **Wafergen iCell8** were derived from Wang et al. 2019 (Table 2)[12]. For the **Wafergen iCell8** it was assumed that 8 samples (one per dispensing nozzle) can be processed on one chip in parallel, thus decreasing the costs by a factor of 8. The **DisCo** cost estimate includes reagents for library generation, i.e. the costs for beads, oil, reverse-transcription reaction, exonuclease treatments, PCR reaction, and library preparation (Nextera XT).

**Physical setup**

Chips were mounted on an IX51 inverted microscope (Olympus). Each chip was monitored with an XiC (Ximea, MC031MG-SY-UB) camera, interfaced with a computer with the following specifications: Windows 10 Enterprise (Microsoft) operating system, Ryzen Threadripper 1950X processor (AMD), 32 GB RAM memory. Solenoid valves were controlled via the NI USB-6501 controller (National Instruments). The output signals from the controller were amplified with a ULN2803 IC (Texas Instruments), and connected to solenoid valves (Festo, MHA1-M1H-3/2O-0,6-HC). An OB1 Mk3 pressure controller (Elveflow) was used for proportional pressure regulation.

**Machine-vision software**

The software for cell detection and coordination was implemented in C++. Camera images were obtained with the XiApi library (version 4.15). Images were processed in real-time using the OpenCV computer vision library (version 3.4). A schematic visualization of the particle detection algorithm is depicted in **Supplementary Figure 1B**. Briefly, a detection ROI was extracted by cropping after which a gaussian blur was applied to the resulting image. Two subsequent images were subtracted, and the resulting image converted to a binary image by intensity thresholding. The binary image was dilated to fill potential holes. Finally, contours were detected using the *findContours* function, and classified for area and circularity. Upon particle detection, the particles were properly positioned by valve oscillation and monitoring of the ROI at the target zone (**Supplementary Figure 1C**). Once two particles were positioned in their respective target zones, particles were co-ejected by pressurization of the dropleting valve, and the droplet was sheared by actuation of the oil valve.

**Microfluidic chip design and fabrication**

The design of the microfluidic chip for deterministic co-encapsulation is presented in **Supplementary Figure 1A**. Chips were designed using Tanner L-Edit CAD software (Mentor, v 2016.2). 5-inch chromium masks were exposed in a VPG200 laser writer (Heidelberg instruments) for both the control and flow layer. Masks were developed using an HMR 900 mask processor (Hamatech). For the control layer, a thick SU8 photoresist layer was deposited with an LSM-200 spin coater (Sawatec), exposed on a MJB4 single side mask aligner (SussMicroTec), and manually developed. The SU8 processing steps were carried out according to manufacturer’s instructions for the 3010 series (Microchem). For the flow layer, wafers were produced using AZ40 XT (Microchem) positive photoresist on the ACS200 coating and developing system (Gen3, SUSS MicroTec). Developed master-wafers were reflowed for 45 - 75 seconds at 120$^{o}$C on a hotplate until channels appeared round under an inspection microscope. The control layer master-wafers were used as molds for PDMS chips after passivation with 1 % silane dissolved in HFE. For the flow layer, master-wafers were used to generate replica molds for chip production. To this end, the primary replica mold was obtained by mixing PDMS:Curing-Agent at 10:1 using a centrifugal mixer (Thinky), degassing for 15 minutes, and curing for 60 minutes at 80°C. The PDMS-based primary replica mold was then sylanized and subsequently used to obtain secondary replica molds utilized for PDMS flow layer production. The PDMS flow layer was fabricated PDMS:Curing-Agent at 5:1, degassed and cured at 80$^{o}$C for 30 minutes. The control layer was fabricated by spin coating PDMS:Curing-Agent at 20:1 on the flow layer waver at 650 rpm for 35 seconds with 15 seconds ramp time followed by baking at 80$^{o}$C for 30 minutes. Cured PDMS was then cut from the flow layer secondary replica mold and flow layer inlet holes were punched with a 0.5 mm diameter biopsy punch. The two PDMS layers were manually aligned and bonded at 80$^{o}$C for at least 60 minutes. Assembled and cured PDMS chips were cut from the molds and control layer inlet holes were punched. Finally, chips were oxygen plasma activated (45 seconds at ~500 mTorr O2) and bonded to a surface activated glass slide followed by incubation at 80$^{o}$C for at least 2 hours. Materials and reagents are listed in the Material and reagent list, point 1.

**Microfluidic device handling**

Prior to use, the microfluidic chip was placed on an inverted microscope and control layer inlets were connected to solenoid valves with water primed tygon tubing. Control layer channels were primed with $dH_2O$ in tygon tubing for ~10 minutes by pressurizing the solenoid valves. If the chip was being used for the first time, cell, bead, and dropleting on-chip valves were equilibrated by oscillation of the corresponding solenoid valves for at least 10 minutes at 2 actuations per second. After priming, the dropleting valve was connected to an OB1 (Elveflow) pressure regulator for proportional actuation. The flow layer was connected the following way: oil, bead, cell inlets and sample outlet to Prot/Elec gel loading tips; waste outlet to tygon tubing terminating in a falcon tube. For inlet pressurization of the Prot/Elec gel loading tip connected inputs, the bead and cell inlets were connected to the OB1 pressure regulator. The oil inlet was continuously pressurized at 1.7 psi. Cell, bead, and oil Prot/Elec tips were filled with cell buffer, bead solution, and oil, respectively. Subsequently, the chip was primed in the following order: 1. cell channel, 2. bead channel, 3. oil channel. After priming, the bead and cell channels were washed for 5 - 10 minutes by running the solutions at low pressure. All priming and washing solutions were directed in the waste outlet. Finally, the sample outlet was primed with oil. Stuffer droplets, containing lysis buffer and RNase inhibitor, were generated on a Drop-seq chip[14] and added on top of the oil-primed sample outlet tip without introducing air bubbles. Materials and reagents are listed in the Material and reagent list, point 2.

**cDNA generation and library preparation**

After bead-cell in droplet co-encapsulation, the gel loading tip containing the sample droplets were transferred to a bead collection chip inlet[14] (cp-chip). Droplets in the tip were flushed to a bead collection chip. Subsequent to bead capture, washing was performed as in the Drop-seq protocol with SSC and reverse transcription buffer directly on the cp-chip. Reverse transcription solution was added to the beads in the recovery chip, and the recovery chip was placed on a heating block to perform first strand cDNA synthesis (RT) for 90 minutes at 42°C. After the RT reaction, beads were washed on the recovery chip with TE-SDS once, with TE-TW twice, and with Tris once. The beads were treated with Exonuclease I for 45 minutes at 37°C to remove single-stranded oligonucleotides on the beads. After Exonuclease I treatment, beads were washed with TE-SDS once, with TE-TW twice (as after RT). Beads were then eluted from the recovery chip in $dH_2O$. cDNA was amplified for 18 – 23 cycles using Kapa HiFi Hot start ready mix. cDNA was purified with CleanPCR magnetic beads (0.6X ratio) to remove small cDNA fragments and primers. The cDNA concentration was measured using Qubit, and cDNA quality was assessed using a Fragment Analyzer (Agilent). cDNA was tagmented with in-house Tn5[15] for 6 minutes at 55°C. Next, the reaction was stopped with SDS and the tagmented library was amplified for 15 cycles using Kapa HiFi kit. Libraries were then purified using CleanPCR magnetic beads (0.6X ratio) and quantified using Qubit HS kit and Fragment analyzer (Agilent). Finally, size-selected and purified libraries were sequenced on a NextSeq 500 system (Illumina) following recommendations from the original Drop-seq protocol (20 bp for read 1 and 50 bp for read2)[16]. Material and reagents are listed in the Material and reagent list, points 3 - 10.

**Mammalian cell culture handling for species mixing experiment**

For benchmarking the DisCo platform, HEK 293T (ATCC Cat. No. SD-3515) and murine brown preadipocyte cells (iBA; provided by Prof. Christian Wolfrum's laboratory, ETH Zürich) were used. Cells were cultured to 90% confluency in Glutamax DMEM supplemented with FBS and penicillin-streptomycin. Prior to use, cells were washed with PBS, dissociated with Trypsin-EDTA, washed with cell wash buffer and counted with Trypan blue live-dead stain using a Countess cell counter (Invitrogen). Cells were mixed in a 1:1 ratio, adjusted to 20 cells/µL, re-suspended in cell loading buffer, and finally loaded on the DisCo chip. Material and reagents are listed in the Material and reagent list, point 11.

**Droplet content and co-encapsulation performance quantification**

As for conventional DisCo runs, experiments were set up with Chemgen beads and varying concentrations of HEK 293T cells. Approximately 100 co-encapsulations were performed and recorded. The recorded video data was manually reviewed and droplet contents and passing cells and beads counted (**Figure 1F**).

**Benchmarking DisCo efficiency using the DISPENCELL platform**

To benchmark single-cell recovery efficiencies throughout the complete DisCo workflow, we quantified HEK 293T (ATCC Cat. No. SD-3515) cells utilizing the DISPENCELL pipetting robot (SEED Biosciences SA). Prior to use, HEK 293T cells were diluted to 20 cells/µL. Cells were loaded into the DISPENCELL tip and then dispensed directly into a Prot/Elec gel loading tip containing cell loading buffer. Cells were then processed with DisCo and libraries prepared as described above.

**Organoid cell culture and handling**

Isolation of the Lgr5-eGFP$^{+}$ stem cells and initial culture was performed as previously described[17]. For the developmental time-course experiments, organoids were dissociated to single-cells, live Lgr5$^{+}$-eGFP cells isolated using a FACS ARIA II (BD) and embedded in Matrigel. After Matrigel polymerization, cells were cultured in ENR CV medium supplemented with thiazovivin ROCK inhibitor.

Growth factors (E, N, R, C, V) were replenished after 2 days of culture. At Day 3 of culture, a full medium change was performed to differentiation growth medium (ENR only). At Day 5, growth-factors (E, N, R) were replenished. Organoids were sampled at Day 3 (S0), prior to the medium change, at Day 4 (S1), at Day 5 (S2), and at Day 6 (S3).

Single organoids were collected by dissolving Matrigel with ice-cold Cell Recovery Solution for approximately 5 minutes, while carefully pipetting up and down with a 1000 µL pipette. Subsequently, single organoids were isolated by hand-picking after which they were transferred to a Nunc microwell culture plate with single organoid dissociation mix. Single organoids were dissociated by combining trituration using siliconized pipette tips every 5 minutes and incubation at 37°C for 15 minutes. Following dissociation, cell suspensions were diluted in cell loading buffer in the loading tip connected to the DisCo chip. Materials and reagents are listed in the Material and reagent list, points 12 - 16.

**RNA Fluorescence in situ hybridization (RNAscope) on intestinal organoids**

For the RNAscope assay, organoids in matrigel were fixed in 4% PFA at 4°C overnight. The next day, organoids were washed with PBS and embedded in histogel. Histogel blocks were subsequently infiltrated with paraffin using a standard histological procedure (VIP6, Sakura). RNAscope Multiplex Fluorescent V2 assay was performed according to the manufacturer's protocol on 4 µm paraffin sections, hybridized with the probes Mm-Ly6a-C2, Mm-Fabp1-C1, Mm-Muc2-C2, Mm-PpiB-C2 positive control, and Duplex negative control at 40°C for 2 hours and revealed with TSA Opal650 for C1 channel and TSA Opal570 for C2 channel. Tissues were counterstained with DAPI and mounted with Prolong Diamond Antifade Mountant. Slides were imaged on an Olympus VS120 whole slide scanner (Olympus). The resulting images were converted to the TIFF file format using the Fiji (version 1.52p) plugin BIOP VSI Reader (version 7). ROIs were extracted using a custom Python (version 2.7.15) script and the PIL library (version 6.2.2). Brightness of the extracted ROIs was adjusted in Fiji: Images of one target were loaded, stacked, brightness adjusted for the whole stack using the *setMinAndMax()* function. Finally, images were unstacked, merged with other channels, and exported as PNG files. Materials and reagents are listed in the Material and reagent list, points 17 - 18.

**Sequencing, analysis, barcode correction**

The data analysis was performed using the Drop-seq tools package (version 2.3.0, https://github.com/broadinstitute/Drop-seq/releases/tag/v2.3.0)[3,16] on the EPFL SCITAS HPC platform. After trimming and sequence tagging, reads were aligned to the human (hg38), mouse (GRCm38), or mixed reference genomes[3] (GSE63269), depending on the origin of the cellular input material, using STAR (version 2.7.0.e)[18]. Following alignment, BAM files were processed to obtain initial read-count matrices (RCM) per sample (Note: DGE summary files were used for experiments displayed in **Figure 1H** and **Figure 1I**). Cell barcodes were prefiltered at > 35 UMIs (for the species mixing experiment, the sum of 35 UMIs for both species was used as a prefiltering criterion). Graphs were built by identifying barcodes connected by Levenshtein distance 1. For each graph, the barcode containing the highest number of UMIs was identified as the central barcode. The graphs were pruned (barcodes removed) at a Levenshtein distance > 2 to the central barcode, the remaining barcodes in the graph were merged.

For cell recovery efficiency experiments using the DISPENCELL platform (**Figure 1I**) and for Drop-seq comparison experiments (**Figure 1J**) barcodes encompassing at least 500 UMIs were compiled into the RCMs. Additionally, prior to Drop-seq comparison experiments, processed BAM files were down sampled to the same read depth using samtools (http://www.htslib.org/doc/samtools.html). Box plot elements depicting UMI counts per cell (**Figure 1I**) represent the following values: centerline, median; box limits, upper/lower quartiles; whiskers, 1.5x interquartile range; points, UMIs per cell.

**Time course organoid kinetic analysis**

RCMs were further processed via R (version 3.6.2) using *Seurat* (version 3.1.1) and *uwot* (version 0.1.3)[19]. Per individual organoid-RCM cells with > 800 features, < 7.5% mitochondrial reads were retained in the analysis. The time course kinetics of organoids were processed in three independent experiments, which were considered as three individual batches. The three independent experiments were merged using *FindIntegrationAnchors(list(experimental_batches), anchor.features = 80, dims = 1:12, k.filter = 200, k.anchor = 8)* and *IntegrateData().* Data was scaled and PCAs computed using default settings. Uniform Manifold Approximation and Projection (UMAP) dimensional reduction via *RunUMAP()* and *FindNeighbors()* were performed using the first 12 PCA dimensions as input features. *FindClusters()* was computed at resolution 0.75. Merged data was visualized using the Seurat intrinsic functions *VlnPlot()*, *FeaturePlot()*, *DotPlot(), DimPlot().* Differentially expressed genes per cluster were identified using *FindAllMarkers()* using default parameters. The Seurat-Object is accessible via GSE148093*.* Cumulative Z-scores were calculated based on the scaled expression per cell across the defined gene signatures[20,21]. Pie-chart, bubble-plot and bargraph visualizations were carried out with *ggplot2*.

**Slingshot analysis**

The trajectories were constructed using the Slingshot wrapper implemented in the dyno package (https://github.com/dynverse/dyno)[22]. The method was provided with the first 5 dimensions of a multi-dimensional scaling as dimensionality reduction, the clustering as described earlier, and the stem cell cluster as starting cell population. All other parameters were left at default settings. Genes that change along the trajectory were ranked using the *calculate_overall_feature_importance* function from the dynfeature package (version 1.0, https://github.com/dynverse/dynfeature), and the top 50 differentially expressed genes were selected. The dynplot package (version 1.1, https://github.com/dynverse/dynplot) was used to plot the trajectory within a scatterplot and heatmap.

**Psupertime analysis**

Cell labels and sample-day labels were extracted from the merged and batch-corrected meta-data of the Seurat object to run *psupertime,* a method of identifying genes relevant to biological processes using cell-level temporal labels to build a l1 regularised ordinal logistic regression model (Macnair & Claassen, biorxiv 2019)[23]. Sample-day labels indicating the experimental temporal order were used to conduct a *psupertime* analysis on batch-corrected and normalized gene expression data of cells, with selected cell type labels. The analysis was performed including all genes and encompassing a 10-fold cross-validation using default settings. Genes with coefficients (beta-values) greater than zero were considered relevant for the temporal expression dynamics. Expression of relevant genes was plotted per organoid per cell.

**Material and reagent list for all experiments**

*Material information is listed in the following format: Material name (vendor, ordering number). Reagent information is listed in the following format: Reagent name (final concentration in the solution, vendor, order number).*

1. For microfluidic device fabrication SU8 3010 (Microchem) negative photoresist, AZ40XT (Microchem) positive photoresist, HFE-7500 (3M, Novec 297730-93-9), Trichloro(1H, 1H, 2H, 2H - perfluorooctyl) silane (1%, Aldrich, 448931), and biopsy punchers (Darwin microfluidics, KPUNCH05) were used.
2. For microfluidic device handling Prot/Elec 200 µL gel loading tips (Biorad, #223-9915), $dH_2O$ (Invitrogen, 10977035), tygon tubing (Cole Palmer, GZ-06420-02), beads (Chemgenes, lot 051917, Macosko-2011-10), droplet generation oil (Biorad, 186-4006), murine RNase inhibitor (100 U, NEB, M0314L) were used. Cell wash buffer was prepared using PBS (1X, Gibco, 14190-094) and BSA (0.01%, Sigma, B8667). Cell loading buffer was prepared using PBS (1X, Gibco, 14190-094), Optiprep (6%, Sigma, D1556), and BSA (0.01%, Sigma, B8667). Lysis buffer was prepared from Optiprep (28%, Sigma, D1556), Sarkosyl (2.2%, Sigma, L7414), EDTA (20 mM, Sigma, 3690), Tris (100 mM, Sigma, T2944), DTT (50 mM, Applichem, A2948,0005).
3. For sample washing prior to reverse transcription, SSC (6X, Sigma, S6639) and $dH_2O$ (Invitrogen 10977-035) were used.
4. For reverse transcription (RT) reaction $dH_2O$ (Invitrogen, 10977-035), Ficoll PM-400 (4%, Sigma, F5415), dNTPs (1mM, Thermo, R0193), murine RNase inhibitor (100U, NEB, M0314L), Maxima H- reverse transcriptase (500 U, Thermo Scientific, EP0753), Template Switching Oligo (AAGCAGTGGTATCAACGCAGAGTGAATrGrGrG, 2.5 µM, IDT) were used in a total volume 50 µL per reaction.
5. For exonuclease I reaction exonuclease I (100 U, NEB, M0293L) and exonuclease buffer were used in a total volume 50 µL per reaction.
6. For cDNA amplification Kapa HiFi Hot start ready mix 2X (Roche, KK2602), $dH_2O$ (Invitrogen, 10977035), and SMART PCR primer (AAGCAGTGGTATCAACGCAGAGT, 0.8 µM, IDT) used in a

total volume 50 µL per reaction. CleanPCR magnetic beads (0.6X ratio, GC biotech, CPCR-0050), Fragment Analyzer (Agilent, DNF-474-0500 kit), and Qubit HS sensitivity kit (Invitrogen, Q33231) were used for cDNA purification and quantification.

7. For library preparation in-house produced Tn5 was used. To stop tagmentation, SDS was used (0.2%, Sigma, 71736). For library amplification Kapa HiFi kit with dNTPs (Roche, KK2102), P5 SMART PCR (AATGATACGGCGACCACCGAGATCTACACGCCTGTCCGCGGAAGCAGTGGTATCAA CGCAGAGT*A*C, 0.3 µM, IDT), custom Nextera oligos[24] (0.3 µM, IDT) and $dH_2O$ (Invitrogen, 10977035) were used. Libraries were purified and quantified using CleanPCR magnetic beads (0.6X ratio, GC biotech, CPCR-0050), Fragment Analyzer (Agilent, DNF-474-0500 kit), and Qubit HS sensitivity kit (Invitrogen, Q33231).
8. TE-TW wash buffer was prepared in $dH_2O$ (Invitrogen, 10977035) using Tris (10 mM, Sigma T2944), EDTA (1mM, Sigma, 3690), and Tween 20 (0.01%, Sigma, P9416).
9. TE-SDS wash buffer was prepared in $dH_2O$ (Invitrogen, 10977035) using Tris (10 mM, Sigma, T2944), EDTA (1 mM, Sigma, 03690), and SDS (0.5%, Sigma, 71736).
10. Tris wash buffer was prepared in $dH_2O$ (Invitrogen, 10977035) using Tris (10 mM, Sigma, T2944).
11. For mammalian cell culture dissociation and counting Trypsin-EDTA (Gibco, 25200056) and trypan blue were used (0.4%, Thermo Fisher Scientific, T10282). Cell culture medium was prepared using DMEM Glutamax (Gibco, 10565018), FBS (10%, Gibco, 10270106) and penicillin-streptomycin (100 U/mL, Gibco, 15140122). Cell wash and cell loading buffers were prepared as described above.
12. Intestinal organoids were cultured in Matrigel (Corning, 356230) with organoid base medium (described in point 13) supplemented with ENR (+ CV where indicated) and rock inhibitor (where indicated, Sigma, Y0503).
13. Organoid base medium was prepared using DMEM/F12 (Gibco, 11320033), Hepes (100 mM, Gibco, 15630056), penicillin-streptomycin (100 U/mL, Gibco, 15140122), B27 supplement (1 µM, Gibco, 17504-044), N2 supplement (1 µM, Gibco, 17502001), and N-Acetyl-L-cysteine (1 µM, Sigma, A9165).

14. ENR medium was prepared using base medium (as above), EGF (E, 50 ng/mL, LifeTechnologies, PMG8043), mNoggin (N, 100 ng/mL, produced in-house), R-spondin (R, 1 µg/mL, produced in-house).
15. ENR CV medium was prepared with addition of CHIR (C, 3 µM, CalBiochem, CHIR99021), and Valproic acid (V, 3 mM, Sigma P4543) to ENR medium.
16. Single-organoid single-cell dissociation mix was prepared using PBS (Gibco, 14190-094), *B. licheniformis* protease (10 mg/mL, Sigma P5380), EDTA (5 mM, Sigma 03690), EGTA (5 mM, BioWorld, 40520008-1), DNase I (10 µg/mL, Roche 11 284 932 001), and Accutase (0.68X, Sigma, A6964) in a total volume 20 µL per reaction. For single organoid dissociation Nunc MicroWell plates (Nunc, 438733) and siliconized p10 pipette tips (VWR, 53509-134) were used.
17. For intestinal organoid preparation for RNAscope, cold Cell Recovery Solution (Corning, 354253), Histogel (Thermo Scientific, HG-4000-012), Paraformaldehyde (4%, PFA, Electron Microscopy Sciences, 15714) were used.
18. For the RNAscope assay, organoids were stained using RNAscope Multiplex Fluorescent V2 assay (ACD Bio-Techne, 323110), Ly6a probe (ACD Bio-Techne, 427571-C2), Fabp1 probe (ACD Bio-Techne, 562831), Muc2 probe (ACD Bio-Techne, 315451-C2), PpiB probe (ACD Bio-Techne, 313911-C2), Duplex negative control (ACD Bio-Techne, 320751), TSA Opal650 (Perkin Elmer, FP1496001KT), TSA Opal570 (Perkin Elmer, FP1488001KT), and Prolong Diamond Antifade Mountant (Thermo Fisher, P36965).

**Public data distribution**

The GEO accession number for scRNA-seq data reported in this paper is GSE148093. For reviewing purposes, a temporary access token is: ejwxsgekplwdzwv .

### 2. Supplementary Figures

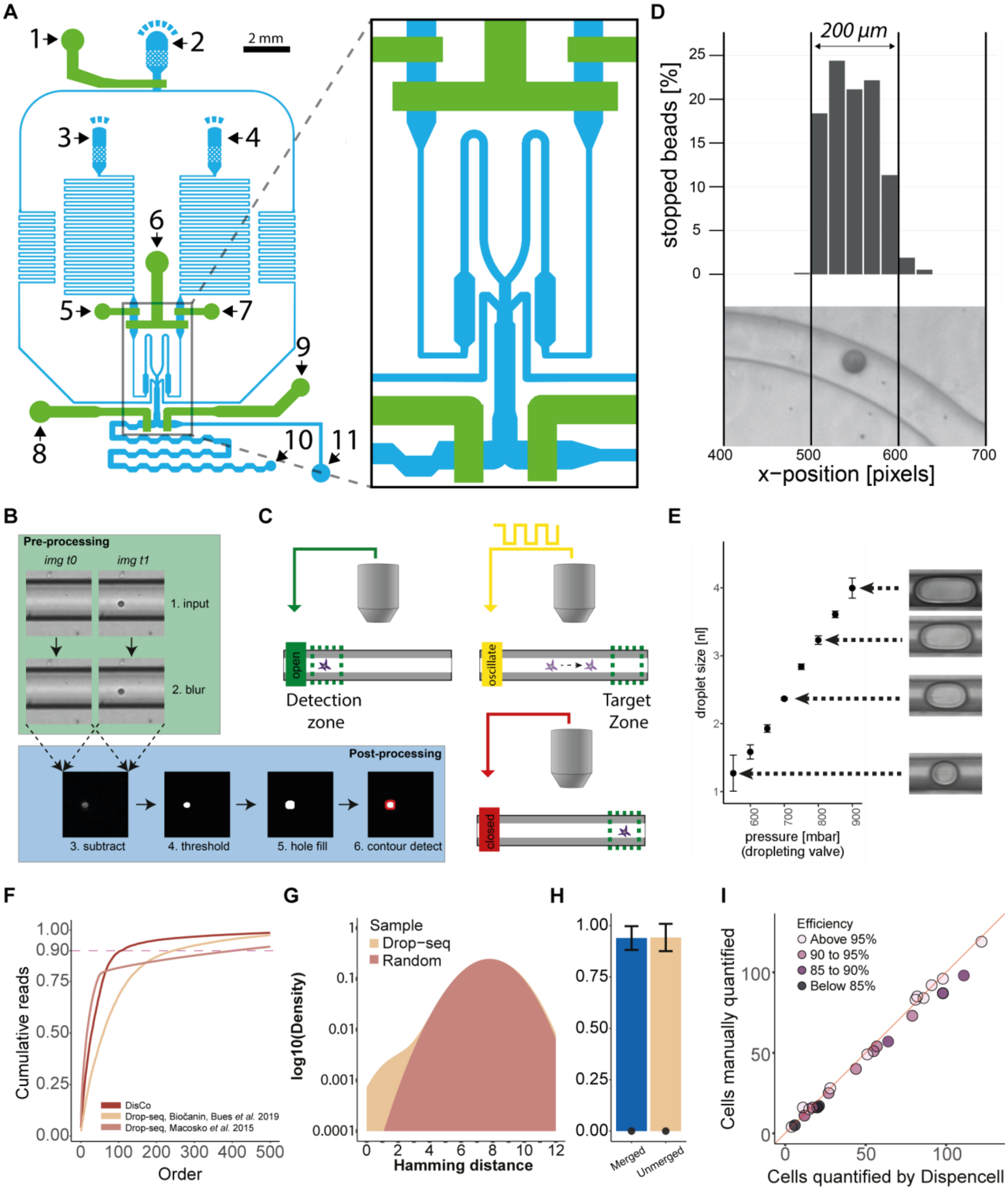


**Supplementary Figure 1:** (**A**) Schematic of the DisCo device design (blue: flow layer, green control layer). 1: oil valve, 2. oil inlet, 3. cell inlet, 4. bead inlet, 5. cell valve, 6. dropleting valve, 7. bead valve, 8. sample valve, 9. waste valve, 10. sample outlet, 11. waste outlet. (**B**) Real-time image processing for particle detection. Two consecutive images are despeckled by gaussian blurring, and subtracted. The resulting image is thresholded and holes are filled by dilatation. Finally, contours are detected and classified by size and circularity thresholding. (**C**) Particle positioning by valve oscillation. Approaching particles are detected in the detection zone. Once a particle is detected, the channel valve is oscillated to induce discrete movements of particles. Oscillation is terminated once correct placement of a particle is achieved. (**D**) Stopping accuracy in a defined window. Beads (n = 744) were positioned using valve oscillation, their position was manually determined within the stopping area. Scale was approximated from channel width. (**E**) Volume-defined droplet on-demand generation by valve pressurization. Droplets (n = 68, ~8 per condition) were produced by pressurizing the dropleting valve at different pressures. Size was determined by imaging the dropleting process. Volumes were calculated from the imaging data based on droplet length and channel geometry. Thus, they should be considered an approximation. Error bars represent standard deviation. The channel width of displayed images is 250 µm. (**F**) Cumulative reads per barcode (n = 500) for DisCo and two Drop-seq experiments[3,14]. (**G**) Hamming distances between all 12 nt barcodes of a Drop-seq experiment, and generated 12 nt random barcode sequences representing the probability density for each set of barcodes. (**H**) Species purity (bars) and doublet ratio (dots) for unmerged and merged barcodes. Error bars represent standard deviation. (**I**) Correlation of the number of manually counted cells by fluorescence microscopy and the number of cells quantified by the DISPENCELL platform.

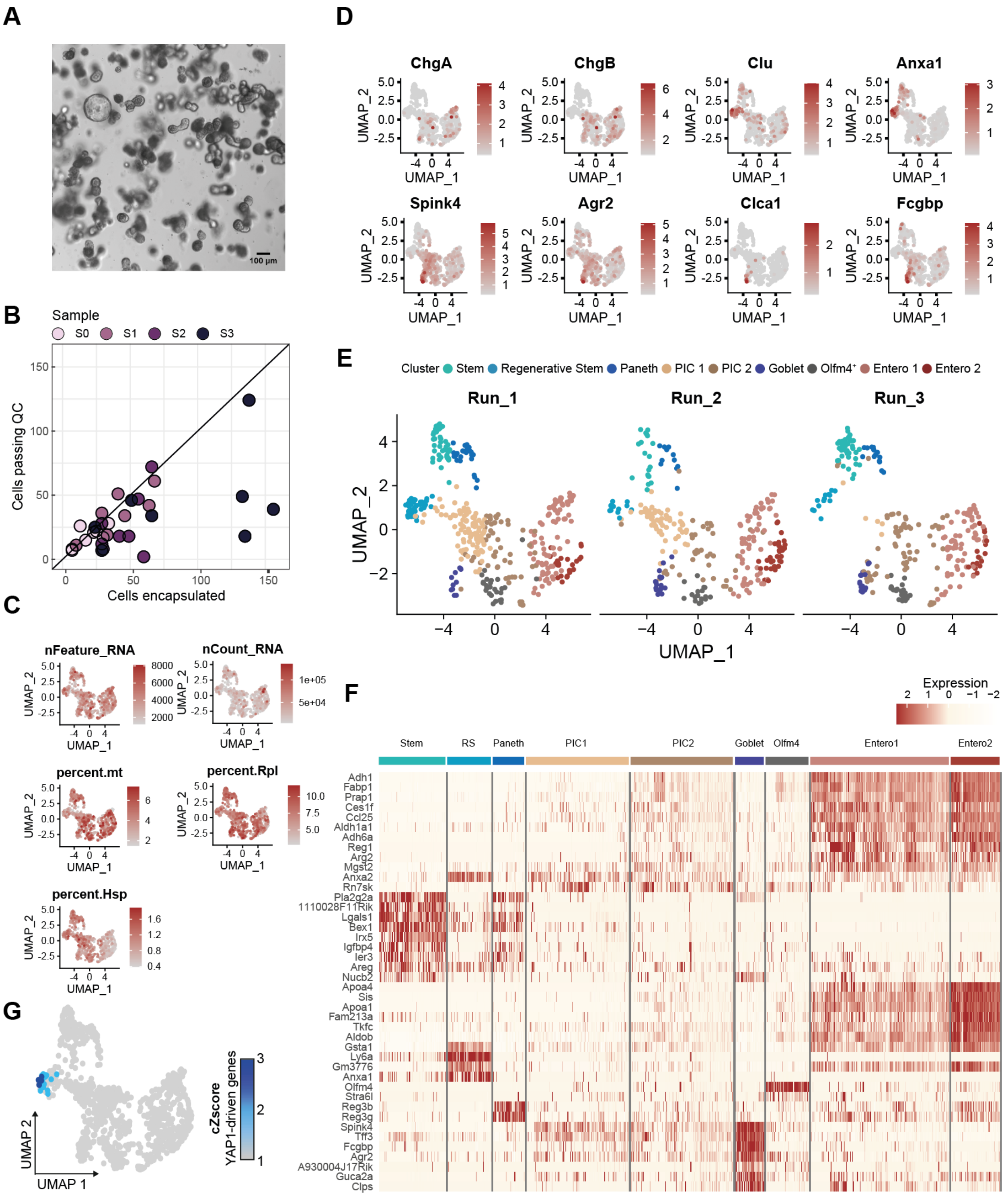


**Supplementary Figure 2: (A)** Representative brightfield image of a differentiated organoid culture from single LGR5⁺ cells. (**B**) Correlation of encapsulated cells on-chip with the number of cells detected after sequencing (Cells passing QC, filtered above 800 genes/cell). (**C**) UMAP embedding colored by number of detected UMIs per cell, number of detected genes per cell, percentage of mitochondrial reads, and percentage of reads mapping to genes coding for respectively ribosomal proteins (Rpl), and heat-shock proteins (Hsp). (**D**) UMAP embedding colored by expression of selected marker genes (*Clu*, *Anxa1*, *Spink4*, *ChgB*, *ChgA*, *Agr2*, *Clca1*, and *Fcgbp*). (**E**) UMAP embedding for each of the three independent experimental batches colored by cluster annotation. (**F**) Heatmap of top DE genes per annotated cluster. (**G**) YAP1 target gene activity on UMAP embedding. The expression of genes that are positively regulated by YAP1[21] was calculated as the cumulative Z-score and projected on the UMAP embedding of all sequenced cells.

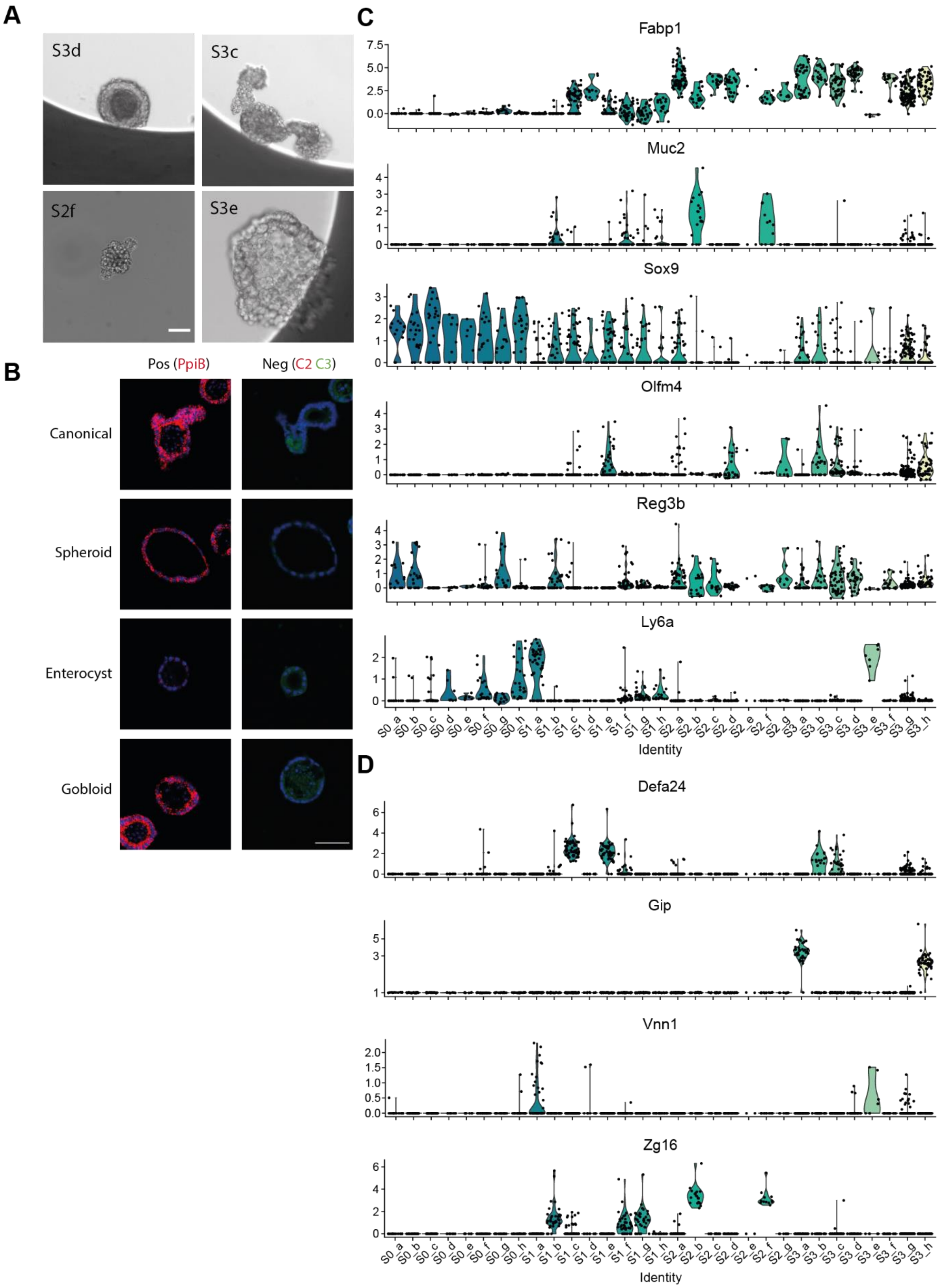
A
S3d
S3c
S2f
S3e
B
Pos (PpiB)
Neg (C2 C3)
Canonical
Spheroid
Enterocyst
Gobloid
C
Fabp1
Muc2
Sox9
Olfm4
Reg3b
Ly6a
Identity
D
Defa24
Gip
Vnn1
Zg16
Identity

**Supplementary Figure 3:** (**A**) Selected organoids imaged in microwell plates before dissociation to single cells. Scale bar 50 µm. (**B**) RNAscope controls for organoids shown in **Figure 3C**. Positive control (*PpiB*), and negative control (Duplex negative). Scale bar 50 µm. (**C**) Violin plots showing marker gene expression (*Fabp1, Muc2, Olfm4, Sox9, Reg3b, Ly6a*) per organoid. (**D**) Violin plot showing the expression of selected genes (*Defa24, Gip, Vnn1, Zg16*) identified via psupertime analysis per individual organoid.

### 3. Supplementary Tables

**Supplementary Table 1**: (**Left Subtable**) Performance summary of established scRNA-seq platform technologies. Performance metrics were derived from literature. Noteworthy, as for lack of consensus experiments, metrics represent different values. (References and calculation of metrics are detailed in the Material and Methods section). (**Right Subtable**) Performance metrics calculated for the DisCo system are presented in this study.

| **Approach** | **Droplets (stochastic)** | | | **FACS & plate based** | | **Traps** | **Microwells** | | | **Droplets (deterministic)** |
|---|---|---|---|---|---|---|---|---|---|---|
| **Technology** | 10X Chromium | inDrop | Drop-seq | Smart-seq2 | Cel-seq2 | Fluidigm C1 | iCell 8 | Seq-well | | DisCo (this study) |
| **Min input** | 500 | 1,000 | 50,000 | 10,000 | 10,000 | 200 | 1,600 | 400 | | < 50 |
| **Efficiency** | 45%* | 25%* | 2.3%* | - | - | 45%** | 43%** | 30%* | | 75%* |
| **$/cell (100 output cells)** | $20 | $2.1 | $6 | $10.6 | $3.6 | $29 (96 cells) | $5 | $2.2 | | $1 |
| **$/cell (100 input cells)** | $44.4 | $8.4 | $260.9 | - | - | $62.2 | $11.6 | $7.5 | | $1.3 |
| **Additional remarks or limitations** | Multiplexing possible, yet requires multiple washing procedures[25,26]. Substantial efficiency losses expected. | | | Fluorescent labeling necessary<br>Expensive to scale up (automation) | | Size-selective properties [8,27] | High initial acquisition cost | | | |

Efficiency estimates: * including cell capture efficiency; ** excluding cell capture efficiency

**Supplementary Table 2:** DE genes for cell clusters

| gene | p_val | avg_logFC | pct.1 | pct.2 | p_val_adj | cluster |
|---|---|---|---|---|---|---|
| **Adh1** | 2.27631673711701e-67 | 1.17602093757581 | 0.991 | 0.639 | 1.82105338969361e-65 | Entero1 |
| **Fabp1** | 8.97391864188553e-65 | 1.03782365836832 | 1 | 0.661 | 7.17913491350843e-63 | Entero1 |
| **Apoa1** | 1.77992432836514e-63 | 0.805522082558538 | 0.981 | 0.557 | 1.42393946269211e-61 | Entero1 |
| **Aldob** | 6.97360352458542e-60 | 0.694678513467298 | 1 | 0.782 | 5.57888281966834e-58 | Entero1 |
| **Prap1** | 2.89992546219975e-59 | 0.819884545216953 | 0.995 | 0.752 | 2.3199403697598e-57 | Entero1 |
| **Ces1f** | 3.67319503250653e-58 | 0.963767358595559 | 0.935 | 0.527 | 2.93855602600522e-56 | Entero1 |
| **Ccl25** | 1.25004133755717e-55 | 0.901771305101302 | 0.986 | 0.693 | 1.00003307004574e-53 | Entero1 |
| **Sis** | 1.20301185332912e-52 | 0.677799265079825 | 0.949 | 0.521 | 9.62409482663297e-51 | Entero1 |
| **Aldh1a1** | 2.70977257321893e-50 | 0.885229974327549 | 0.94 | 0.656 | 2.16781805857514e-48 | Entero1 |
| **Adh6a** | 1.98840147831558e-49 | 0.861278352992536 | 0.875 | 0.412 | 1.59072118265247e-47 | Entero1 |
| **Reg1** | 9.99062930782402e-45 | 1.56303848520865 | 0.847 | 0.494 | 7.99250344625922e-43 | Entero1 |
| **Gsta3** | 3.89871251249365e-44 | 0.805279610297088 | 0.889 | 0.528 | 3.11897000999492e-42 | Entero1 |
| **Fabp2** | 1.29229051817192e-42 | 0.729193895321954 | 0.963 | 0.745 | 1.03383241453754e-40 | Entero1 |
| **Spink1** | 3.66097718899745e-42 | 0.629779702665184 | 0.87 | 0.454 | 2.92878175119796e-40 | Entero1 |
| **Arg2** | 7.98376218566301e-40 | 0.831847381415513 | 0.815 | 0.455 | 6.38700974853041e-38 | Entero1 |
| **Apoa4** | 1.02016690530616e-39 | 0.61007480062237 | 0.838 | 0.442 | 8.16133524244929e-38 | Entero1 |
| **Mgst2** | 7.10101680893117e-39 | 0.851374862110365 | 0.912 | 0.709 | 5.68081344714493e-37 | Entero1 |
| **S100g** | 2.96870148861901e-37 | 0.67331354650757 | 0.894 | 0.539 | 2.37496119089521e-35 | Entero1 |
| **Gsta1** | 1.35306424273308e-32 | 0.472486977624749 | 0.995 | 0.796 | 1.08245139418646e-30 | Entero1 |
| **Tkfc** | 3.50611905883385e-32 | 0.411753509964946 | 0.94 | 0.687 | 2.80489524706708e-30 | Entero1 |
| **Cyp4f14** | 5.38663195657445e-31 | 0.519981900345935 | 0.852 | 0.55 | 4.30930556525956e-29 | Entero1 |
| **Ces2a** | 2.32096694803394e-30 | 0.625871228915386 | 0.991 | 0.83 | 1.85677355842715e-28 | Entero1 |
| **Ugt2b5** | 1.16317697151327e-29 | 0.65462290216724 | 0.759 | 0.446 | 9.30541577210619e-28 | Entero1 |
| **Cideb** | 1.66522526047928e-29 | 0.668807931062889 | 0.838 | 0.55 | 1.33218020838342e-27 | Entero1 |
| **Khk** | 2.3562753913959e-29 | 0.559103120929781 | 0.912 | 0.691 | 1.88502031311672e-27 | Entero1 |
| **Gm3776** | 6.14199666813945e-29 | 0.494460207582878 | 0.889 | 0.593 | 4.91359733451156e-27 | Entero1 |
| **Ces1d** | 1.38677645463885e-28 | 0.400811817763043 | 0.653 | 0.28 | 1.10942116371108e-26 | Entero1 |
| **Ndrg1** | 2.24776560017418e-27 | 0.521412321460839 | 0.87 | 0.601 | 1.79821248013934e-25 | Entero1 |
| **Cyp3a11** | 2.06247550022557e-26 | 0.527798351284679 | 0.736 | 0.384 | 1.64998040018045e-24 | Entero1 |
| **Cyp2b10** | 2.15539125536087e-25 | 0.697335344923729 | 0.704 | 0.387 | 1.7243130042887e-23 | Entero1 |
| **Fam213a** | 6.10640077092692e-25 | 0.301752190721072 | 0.861 | 0.608 | 4.88512061674153e-23 | Entero1 |
| **Cyp2c66** | 1.35874414549653e-23 | 0.488752146117802 | 0.736 | 0.512 | 1.08699531639722e-21 | Entero1 |
| **Gstm4** | 1.47764282293861e-23 | 0.653896742719388 | 0.676 | 0.399 | 1.18211425835089e-21 | Entero1 |
| **Ephx1** | 3.22919737315709e-20 | 0.456673877333623 | 0.806 | 0.531 | 2.58335789852567e-18 | Entero1 |
| **Chp2** | 1.89761043469546e-19 | 0.411199981170197 | 0.764 | 0.508 | 1.51808834775637e-17 | Entero1 |
| **Guca2b** | 6.3835695365107e-19 | 0.432207103895791 | 0.694 | 0.385 | 5.10685562920856e-17 | Entero1 |
| **Acaa1b** | 2.25648025784849e-18 | 0.495888699080675 | 0.634 | 0.364 | 1.80518420627879e-16 | Entero1 |
| **Cyp2c29** | 5.3831327465173e-18 | 0.394391182028341 | 0.597 | 0.299 | 4.30650619721384e-16 | Entero1 |
| **Leap2** | 9.71064455701969e-17 | 0.342055623616844 | 0.653 | 0.361 | 7.76851564561575e-15 | Entero1 |
| **Cyp2d26** | 1.33095364656475e-16 | 0.338288348487044 | 0.667 | 0.379 | 1.0647629172518e-14 | Entero1 |

| | | | | | | |
|---|---|---|---|---|---|---|
| **Mogat2** | 1.97346003982513e-16 | 0.58554835926939 | 0.662 | 0.412 | 1.57876803186011e-14 | Entero1 |
| **Apoc3** | 1.7352844842063e-15 | 0.271366484121131 | 0.616 | 0.309 | 1.38822758736504e-13 | Entero1 |
| **Gsta2** | 2.62559594439782e-15 | 0.250857127832048 | 0.69 | 0.435 | 2.10047675551825e-13 | Entero1 |
| **Golgb1** | 1.29123859560501e-13 | 0.438636271560022 | 0.833 | 0.63 | 1.03299087648401e-11 | Entero1 |
| **Cyp3a13** | 1.30217888157126e-13 | 0.394118685929322 | 0.671 | 0.476 | 1.04174310525701e-11 | Entero1 |
| **Mt4** | 2.18300364431703e-08 | 0.345660220432632 | 0.551 | 0.388 | 1.74640291545362e-06 | Entero1 |
| **Anxa2** | 0.000105107960280338 | 0.376798010321461 | 0.762 | 0.761 | 0.00840863682242703 | PIC1 |
| **Rn7sk** | 4.20752227853046e-22 | 0.49105710835518 | 0.919 | 0.651 | 3.36601782282437e-20 | PIC2 |
| **Pla2g2a** | 3.19123918126354e-40 | 1.3249929078319 | 0.913 | 0.564 | 2.55299134501083e-38 | Stem |
| **1110028F11Rik** | 3.66737905051025e-40 | 1.04343585788738 | 0.846 | 0.46 | 2.9339032404082e-38 | Stem |
| **Lgals1** | 7.35974158937406e-40 | 1.2525285566728 | 0.885 | 0.483 | 5.88779327149925e-38 | Stem |
| **Bex1** | 4.07766717099372e-39 | 1.43857975339842 | 0.923 | 0.586 | 3.26213373679497e-37 | Stem |
| **Irx5** | 1.38160802990555e-30 | 1.01446376098122 | 0.788 | 0.529 | 1.10528642392444e-28 | Stem |
| **Igfbp4** | 5.45407120418318e-27 | 0.818091470946171 | 0.817 | 0.558 | 4.36325696334655e-25 | Stem |
| **Ier3** | 8.66238894755805e-21 | 0.902890680630629 | 0.817 | 0.621 | 6.92991115804644e-19 | Stem |
| **Areg** | 1.44349039107372e-16 | 0.995256095963472 | 0.894 | 0.709 | 1.15479231285898e-14 | Stem |
| **Nucb2** | 4.38795737047699e-10 | 0.541151354998672 | 0.731 | 0.583 | 3.51036589638159e-08 | Stem |
| **Apoa4** | 5.93932200196654e-45 | 1.6534454453666 | 1 | 0.491 | 4.75145760157324e-43 | Entero2 |
| **Sis** | 1.10343694212606e-44 | 1.92728989561058 | 1 | 0.586 | 8.82749553700845e-43 | Entero2 |
| **Apoa1** | 1.1201725751272e-44 | 2.15854194457433 | 1 | 0.624 | 8.96138060101757e-43 | Entero2 |
| **Fam213a** | 1.30222226270094e-43 | 1.92083136662646 | 1 | 0.636 | 1.04177781016075e-41 | Entero2 |
| **Tkfc** | 7.58132539855834e-42 | 1.80677470413554 | 0.987 | 0.724 | 6.06506031884668e-40 | Entero2 |
| **Aldob** | 4.37622510619118e-41 | 2.39954242291639 | 1 | 0.817 | 3.50098008495295e-39 | Entero2 |
| **Prap1** | 8.67990158043496e-41 | 1.85693038324859 | 1 | 0.791 | 6.94392126434797e-39 | Entero2 |
| **Spink1** | 4.84903542738247e-40 | 1.48825704587288 | 0.974 | 0.512 | 3.87922834190598e-38 | Entero2 |
| **Khk** | 3.00508823625212e-39 | 1.34034925225687 | 1 | 0.719 | 2.40407058900169e-37 | Entero2 |
| **Fabp1** | 5.99265196134052e-39 | 2.12550630530773 | 1 | 0.716 | 4.79412156907241e-37 | Entero2 |
| **Adh1** | 1.82859672267536e-38 | 1.59803619061465 | 1 | 0.695 | 1.46287737814029e-36 | Entero2 |
| **Gsta2** | 2.72420461425157e-37 | 1.14631403188223 | 0.961 | 0.452 | 2.17936369140126e-35 | Entero2 |
| **Ccl25** | 6.84495230843095e-37 | 1.27948413836236 | 1 | 0.739 | 5.47596184674476e-35 | Entero2 |
| **Ephx1** | 1.52344584782225e-35 | 1.21448812092998 | 0.987 | 0.559 | 1.2187566782578e-33 | Entero2 |
| **Adh6a** | 2.20786040162389e-34 | 1.12917276504955 | 0.987 | 0.476 | 1.76628832129911e-32 | Entero2 |
| **Cyp4f14** | 2.08329178298781e-33 | 1.21375095416519 | 0.987 | 0.587 | 1.66663342639025e-31 | Entero2 |
| **Gsta1** | 2.59753045339481e-33 | 1.49463348002007 | 1 | 0.827 | 2.07802436271585e-31 | Entero2 |
| **S100g** | 3.10615120049419e-33 | 1.12431617027042 | 0.974 | 0.589 | 2.48492096039535e-31 | Entero2 |
| **Cyp2c66** | 5.35653099425473e-33 | 1.15968719761029 | 0.961 | 0.528 | 4.28522479540378e-31 | Entero2 |
| **Chp2** | 1.02267027061363e-32 | 1.14689618487639 | 0.974 | 0.53 | 8.18136216490906e-31 | Entero2 |
| **Cyp2d26** | 1.41893843629184e-32 | 1.03781759437069 | 0.934 | 0.402 | 1.13515074903347e-30 | Entero2 |
| **Ndrg1** | 9.50957333894437e-32 | 1.13147307348477 | 0.961 | 0.636 | 7.60765867115549e-30 | Entero2 |
| **Leap2** | 9.79198061789397e-32 | 1.02395326084673 | 0.882 | 0.388 | 7.83358449431517e-30 | Entero2 |
| **Cyp3a11** | 3.84843836479325e-31 | 1.24933472690356 | 0.895 | 0.427 | 3.0787506918346e-29 | Entero2 |
| **Guca2b** | 3.07316875882886e-30 | 1.01512661269658 | 0.908 | 0.417 | 2.45853500706308e-28 | Entero2 |
| **Gm3776** | 8.43962680052236e-30 | 1.29978848049862 | 0.987 | 0.632 | 6.75170144041789e-28 | Entero2 |

| | | | | | | |
|---|---|---|---|---|---|---|
| **Ugt2b5** | 1.58240974168786e-29 | 1.06991560499757 | 0.921 | 0.482 | 1.26592779335029e-27 | Entero2 |
| **Reg3a** | 6.55267691058194e-29 | 1.35713088197254 | 0.829 | 0.369 | 5.24214152846556e-27 | Entero2 |
| **Apoc3** | 7.76643242226385e-29 | 0.985880738820278 | 0.842 | 0.338 | 6.21314593781108e-27 | Entero2 |
| **Cyp3a13** | 3.55897381144747e-28 | 1.01591555496027 | 0.908 | 0.487 | 2.84717904915798e-26 | Entero2 |
| **Fabp2** | 3.37223145568505e-26 | 0.887488937761581 | 0.987 | 0.778 | 2.69778516454804e-24 | Entero2 |
| **Cideb** | 5.18165254326076e-26 | 0.929781489565813 | 0.974 | 0.585 | 4.14532203460861e-24 | Entero2 |
| **Aldh1a1** | 2.55441426054888e-25 | 0.893818129793693 | 0.961 | 0.7 | 2.0435314084391e-23 | Entero2 |
| **Ces1f** | 5.35862461659871e-22 | 0.851231933945787 | 0.961 | 0.59 | 4.28689969327897e-20 | Entero2 |
| **Gsta3** | 1.30110417219731e-17 | 0.63389982862055 | 0.947 | 0.581 | 1.04088333775785e-15 | Entero2 |
| **Mogat2** | 5.54585767826874e-16 | 0.674414589926087 | 0.789 | 0.441 | 4.436686142615e-14 | Entero2 |
| **Reg1** | 4.54564681142072e-15 | 0.888576028125212 | 0.803 | 0.555 | 3.63651744913658e-13 | Entero2 |
| **Mir22hg** | 1.33018419475433e-14 | 0.366380428724304 | 0.711 | 0.413 | 1.06414735580346e-12 | Entero2 |
| **Arg2** | 3.26751048825669e-13 | 0.594244489315401 | 0.816 | 0.513 | 2.61400839060536e-11 | Entero2 |
| **Reg3b** | 5.40722600169337e-13 | 0.693131482994203 | 0.855 | 0.585 | 4.3257808013547e-11 | Entero2 |
| **Cyp2b10** | 1.12509349975993e-12 | 0.688885456222288 | 0.75 | 0.434 | 9.00074799807947e-11 | Entero2 |
| **Ces2a** | 7.06803557675554e-12 | 0.396131534280798 | 1 | 0.855 | 5.65442846140443e-10 | Entero2 |
| **Reg3g** | 1.30407635700036e-10 | 0.31181532413503 | 0.882 | 0.636 | 1.04326108560029e-08 | Entero2 |
| **Ephx2** | 1.70167582930346e-10 | 0.437834332901168 | 0.684 | 0.42 | 1.36134066344277e-08 | Entero2 |
| **Mgst2** | 2.72676342101281e-09 | 0.49248922756913 | 0.934 | 0.74 | 2.18141073681025e-07 | Entero2 |
| **Acaa1b** | 1.43762853426396e-08 | 0.386561090746002 | 0.684 | 0.403 | 1.15010282741117e-06 | Entero2 |
| **Mt4** | 1.45515610140142e-07 | 0.271929123109072 | 0.632 | 0.407 | 1.16412488112114e-05 | Entero2 |
| **Ly6a** | 2.01002501530632e-36 | 1.80821629971927 | 0.926 | 0.527 | 1.60802001224506e-34 | RS |
| **Anxa2** | 1.02615198797397e-22 | 1.04409160576179 | 0.971 | 0.745 | 8.20921590379172e-21 | RS |
| **Gm3776** | 3.18379144779195e-20 | 0.950402325556938 | 0.956 | 0.637 | 2.54703315823356e-18 | RS |
| **Anxa1** | 9.29314805494896e-19 | 1.29093095938257 | 0.765 | 0.521 | 7.43451844395917e-17 | RS |
| **Gsta1** | 7.71481200331918e-18 | 0.858095925736984 | 1 | 0.829 | 6.17184960265535e-16 | RS |
| **Areg** | 1.07592965296389e-06 | 0.374112922663416 | 0.838 | 0.721 | 8.60743722371114e-05 | RS |
| **Olfm4** | 7.34682088759632e-38 | 2.10864807562475 | 0.97 | 0.558 | 5.87745671007706e-36 | Olfm4 |
| **Stra6l** | 3.32234482841291e-09 | 0.330277790744615 | 0.716 | 0.544 | 2.65787586273033e-07 | Olfm4 |
| **Reg3b** | 6.19284589582374e-21 | 1.79285294751421 | 0.939 | 0.588 | 4.95427671665899e-19 | Paneth |
| **Reg3g** | 4.23085976422072e-20 | 1.61610736104673 | 0.918 | 0.642 | 3.38468781137658e-18 | Paneth |
| **Pla2g2a** | 5.53846330858548e-14 | 1.11076702995807 | 0.837 | 0.589 | 4.43077064686839e-12 | Paneth |
| **Lgals1** | 2.42086721115051e-12 | 0.625986886999084 | 0.816 | 0.511 | 1.9366937689204e-10 | Paneth |
| **Bex1** | 3.87275526447042e-12 | 0.781270009856096 | 0.878 | 0.609 | 3.09820421157633e-10 | Paneth |
| **Ier3** | 3.95525301117695e-10 | 0.577960555043002 | 0.816 | 0.633 | 3.16420240894156e-08 | Paneth |
| **Igfbp4** | 0.000137577038169854 | 0.459617953534454 | 0.653 | 0.583 | 0.0110061630535883 | Paneth |
| **Spink4** | 3.52440643770931e-28 | 2.97630334738867 | 1 | 0.62 | 2.81952515016745e-26 | Goblet |
| **Tff3** | 4.23028954766408e-27 | 2.52408433135767 | 1 | 0.844 | 3.38423163813126e-25 | Goblet |
| **Fcgbp** | 1.58233080983152e-26 | 2.15228915927589 | 0.956 | 0.412 | 1.26586464786522e-24 | Goblet |
| **Agr2** | 4.13763902264459e-22 | 2.22989599251811 | 0.956 | 0.592 | 3.31011121811567e-20 | Goblet |
| **A930004J17Rik** | 7.15243251983664e-16 | 0.270159357631491 | 0.8 | 0.444 | 5.72194601586931e-14 | Goblet |
| **Guca2a** | 3.66076196916109e-15 | 1.5382791422853 | 0.867 | 0.562 | 2.92860957532887e-13 | Goblet |
| **Nucb2** | 4.38484098082501e-15 | 0.84224961248207 | 0.889 | 0.584 | 3.50787278466001e-13 | Goblet |

| | | | | | | |
|---|---|---|---|---|---|---|
| **Clps** | 1.46445135819238e-14 | 1.43210436908701 | 0.822 | 0.486 | 1.1715610865539e-12 | Goblet |
